# Sources of prey availability data alter interpretation of outputs from prey choice null networks

**DOI:** 10.1101/2023.07.25.549927

**Authors:** Jordan P. Cuff, Maximillian P.T.G. Tercel, Fredric M. Windsor, Ben S.J. Hawthorne, Peter A. Hambäck, James R. Bell, William O.C. Symondson, Ian P. Vaughan

**Affiliations:** School of Biosciences, Cardiff University, Museum Avenue, Cardiff, CF10 3AX, UK; Rothamsted Insect Survey, Rothamsted Research, West Common, Harpenden, Hertfordshire, AL5 2JQ, UK; School of Natural and Environmental Sciences, Newcastle University, Newcastle upon Tyne, NE1 7RU, UK; Durrell Wildlife Conservation Trust, Les Augrès Manor, La Profonde Rue, Trinity, Jersey, JE3 5BP, Channel Islands; Department of Ecology, Environment and Plant Sciences, Stockholm University, 106 91 Stockholm, Sweden

**Keywords:** dietary analysis, field sampling, high-throughput sequencing, metabarcoding, network ecology, null modelling, predictive modelling, resource preference

## Abstract

1. Null models provide an invaluable baseline against which to test fundamental ecological hypotheses and highlight patterns in foraging choices that cannot be explained by neutral processes or sampling artefacts. In this way, null models can advance our understanding beyond simplistic dietary descriptions to identify drivers of interactions. This method, however, requires estimates of resource availability, which are generally imperfect representations of highly dynamic systems. Optimising method selection is crucial for study design, but the precise effects of different resource availability data on the efficacy of null models are poorly understood.
2. Using spider-prey networks as a model, we used prey abundance (suction sample) and activity density (sticky trap) data, and combinations of the two, to simulate null networks. We compared null diet composition, network properties (e.g., connectance and nestedness) and deviations of simulations from metabarcoding-based spider dietary data (to ascertain how different prey availability data alter ecological interpretation.
3. Different sampling methods produced different null networks and inferred distinct prey selectivity. Null networks based on prey abundance and combined frequency-of-occurrence data more closely resembled the observed diet composition, and those based on prey abundance, activity density and proportionally combined data generated network properties most like dietary metabarcoding networks.
4. We show that survey method choice impacts all aspects of null network analyses, the precise effects varying between methods but ultimately altering ecological interpretation by increasing disparity in network properties or trophic niches between null and directly constructed networks. Merging datasets can generate more complete prey availability data but is not a panacea because it introduces different biases. The choice of method should reflect the research hypotheses and study system being investigated. Ultimately, survey methods should emulate the foraging mode of the focal predator as closely as possible, informed by the known ecology, natural history and behaviour of the predator.

## Introduction

Trophic interactions are fundamental to evolutionary and ecological processes (Vázquez & Aizen, 2003). The identity and frequency of interactions are determined by resource preferences and choices, and assessing resource choice is crucial in predicting and understanding trophic dynamics, behaviour and ecology more broadly (Cuff, Tercel, Drake, Vaughan, et al., 2022). For example, assessing the structure of these interactions can provide insight into network assembly, functioning and response to perturbation (Allesina et al., 2008). It is, however, difficult to contextualise independent observations of resource choice given the highly system-dependent nature of such patterns (Vázquez & Aizen, 2003).

Null models facilitate testing of ecological hypotheses by comparing observations with null expectations by representing specific mechanisms (e.g., trait-dependent interactions) and generating random data, with various successful applications across ecology and biogeography (Gotelli, 2001; Gotelli & Graves, 1996). Null modelling can reveal when trophic interactions deviate from random by providing a baseline representation of both the frequency and identity of interactions as would be generated by random foraging (Vaughan et al., 2018; Vázquez & Aizen, 2003). Most simply, this approach can assess how trophic interactions relate to prey abundances in which the most abundant prey are likely to be the most commonly consumed (Agustí et al., 2003; Cuff, Tercel, Drake, Vaughan, et al., 2022; Vaughan et al., 2018). This approach can also provide valuable information on predators (e.g., behaviour, preferences, nutritional requirements), prey (e.g., palatability, detectability, defences, escape ability) and the trophic network in which they exist (e.g., network metrics like nestedness and linkage density), provided that input data represent the relative availability of different prey types to the focal predator. Individual-based approaches to null modelling (i.e., generating null data for each individual), by not fixing network properties *a priori*, reduce constraints on null network generation for comparison against observed networks, leading to more realistic and stochastic null networks (Grimm & Berger, 2016). These approaches can highlight network structures that are not generated by neutral mechanisms, or arise as an artefact of sampling methods, by maintaining the characteristics of the observed data (e.g., the degree of each consumer; Blüthgen et al., 2008; Vaughan et al., 2018). The interaction identities and frequencies, and the concomitant network structures, can nevertheless differ greatly depending on the input data used.

If accurately constructed, null models can elucidate the fundamental mechanisms underpinning species interactions. Null model approaches have therefore been used to explore a range of research questions including prey selectivity changes in response to perturbations (Cuff et al., 2021), seasonal variations in prey availability (Gajski et al., 2023; Verschut et al., 2019), host-parasite-parasitoid specialisation (Ramirez et al., 2022), pollinator preferences across different landscapes (Gómez□Martínez et al., 2022), changes in foraging ecology corresponding with weather conditions (Cuff, Windsor, et al., 2023) and plant-invertebrate commensalisms (Cuff, Evans, Porteous, Quiñonez, et al., 2022). Alongside taxonomic units (e.g., species), the nodes in these networks can represent data such as consumer age class (Davies et al., 2022), functional groups (Méndez□Castro et al., 2020) and environmental context (Cuff, Windsor, et al., 2023), increasing the value and applicability of these models. Many of these examples, particularly those concerning plant resources, assess preferences of active consumers for static resources, for which resource availability is relatively straightforward to estimate, but the interpretation is confounded when both consumer and resource are mobile (e.g., predator-prey systems), and little guidance exists regarding resource availability estimation. To avoid biases caused by improper resource availability estimates, it is paramount that the choice of sampling method aims to closely match the prey available to a predator (Table 1).

**Table 1:**
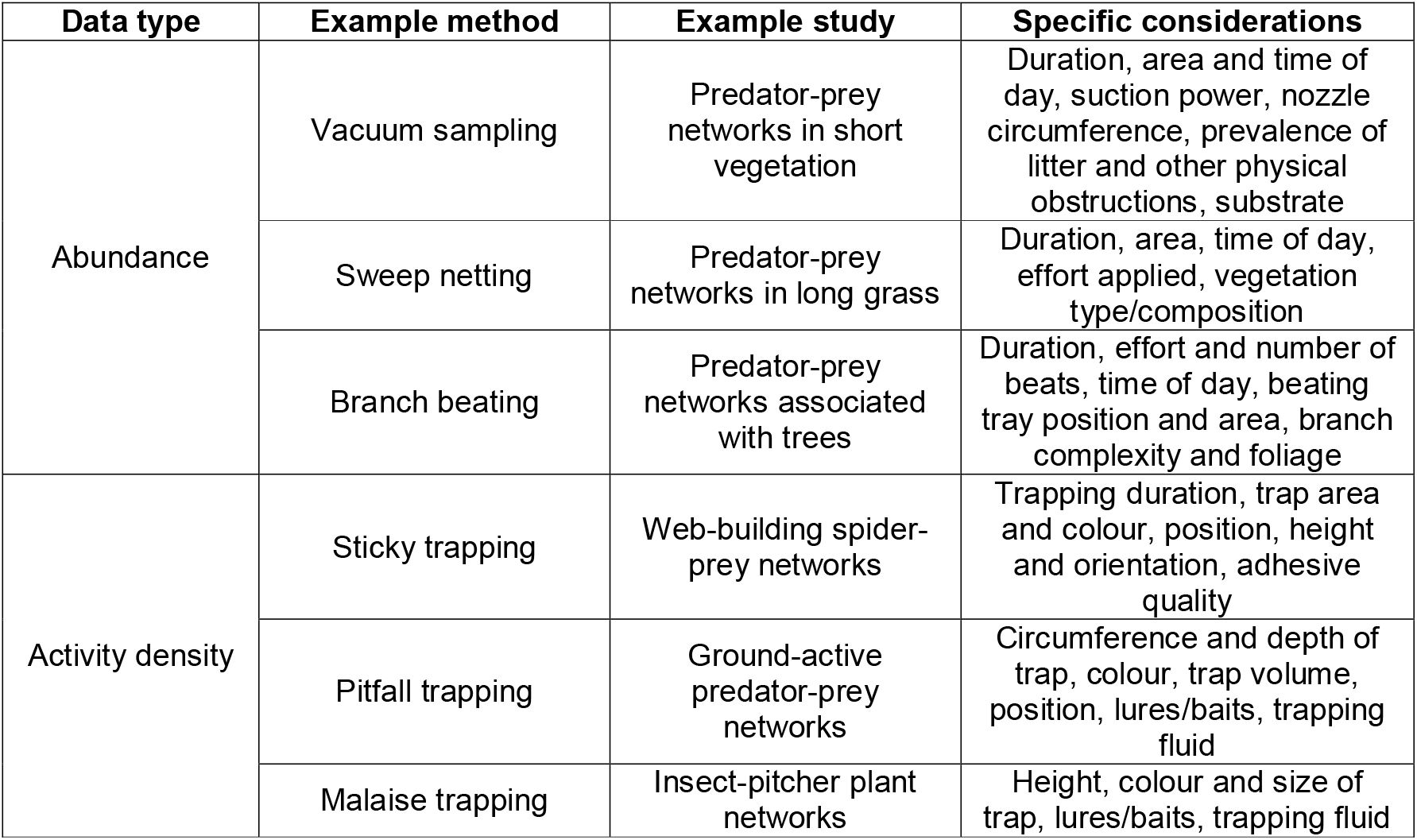
Methods used to generate invertebrate prey availability data, example studies they might be used for and some specific considerations. Methods should be selected to best reflect the experience of focal predators and to reduce eventual biases. Note that all methods may be sensitive to weather conditions.

In this manuscript, we show how survey method choice affects null-model-based resource choice analyses, with significant implications for broader studies relating resource and interaction data to understand drivers of interactions such as predator foraging ecology. Given that spiders can employ both active hunting and sit-and-wait predation, two data types representing prey availability were collected. Prey activity density and abundance samples were collected at each sampling location using sticky traps and suction sampling, respectively. These sampling methods were then used to generate null networks based on dietary data, but we also used two different methods to combine these two estimates into one prey availability index. Using these different prey availability estimates, we tested the following hypotheses: (i) survey method choice affects the results of null model analysis by altering the identity and frequency of simulated trophic interactions, and ultimately network properties; (ii) different measures of prey availability (i.e., abundance and activity density) differ in their relationship to observed interactions, reflecting their emulation of the foraging behaviour of the consumer; and (iii) the type of observed interaction data used alters inferred foraging choices and the structure of null networks. Through discussion of these hypotheses, we also provide guidance for researchers embarking on relevant studies and discuss how to overcome inaccuracies introduced by survey method biases.

## Materials and Methods

### Fieldwork

Data collection was described previously by Cuff, Tercel, et al., (2022). This study pertains to a subset of those data, collected between 1^st^ May and 9^th^ July 2018 at 19 separate locations, for which paired sticky trap and vacuum sample data were collected (described below). Briefly, money spiders (Araneae: Linyphiidae) and wolf spiders (Araneae: Lycosidae) were visually located along transects in two adjacent barley fields at Burdons Farm, Wenvoe in South Wales (51°26’24.8"N, 3°16’17.9"W) and collected from webs and the ground. Transects were randomly distributed across the entire field. Along these transects, separate 4 m^2^ quadrats, at least 10 m apart, were searched and all observed linyphiids and lycosids were collected. Spiders were placed in 100 % ethanol using an aspirator, regularly changing meshing to limit potential cross-contamination. Linyphiids occupying webs were prioritised for collection, but ground-active spiders were also collected. Spiders were taken to Cardiff University, transferred to fresh ethanol and stored at -80 °C in 100 % ethanol until DNA extraction. Extraction, amplification and sequencing of DNA, and bioinformatic analysis is described by Cuff, Tercel, et al., (2022) and Drake et al., (2022), and is also detailed in Supplementary Information 1. The resultant sequencing read counts were converted into relative proportions (all values made to sum to one within each sample) and a mean value across the two primer pairs retained for each taxon within each sample. Relative read abundances were converted to presence-absence data of each detected prey taxon in each individual spider, but relative read abundance data were also retained for separate analyses to compare experimental outcomes between treatments.

To estimate prey availability using sticky traps, we placed one white dry 100 mm x 125 mm trap (Oecos) in the 4 m^2^ quadrat centred at the position where the spider was captured. The trap was suspended with wire approximately 25 mm above the ground to catch falling, crawling and flying invertebrates, and left in place for 72 hours. Invertebrates were identified on the traps under a stereomicroscope. To estimate prey availability using suction sampling, ground and crop stems were sampled using a ‘G-vac’ for approximately 30 seconds at each location. The collected material was emptied into a bag, any organisms immediately killed with ethyl-acetate and material frozen for storage before sorting into 70 % ethanol in the lab. All invertebrates were identified to family level to match the resolution of the least resolved of the metabarcoding-derived trophic interaction data, and due to difficulties associated with identification to finer taxonomic resolution for many taxa. Exceptions included springtails of the superfamily Sminthuroidea (Sminthuridae and Bourletiellidae were often indistinguishable following suction sampling and preservation due to the fine features necessary to distinguish them) which were left at super-family, mites (many of which were immature or in poor condition) which were identified to order level, and wasps of the superfamily Ichneumonoidea which were identified no further due to obscurity of wing venation due to damage following suction sampling.

### Statistical Analysis

All analyses were conducted in R v4.0.3 (R Core Team, 2021) and carried out on invertebrate data at the family or superfamily level. Alongside the dietary data derived from metabarcoding, and prey availability as determined directly by suction sampling (abundance) and sticky trapping (activity density), three additional datasets were generated where two were designed to combine data from the two trapping methods (Figure 1). The first approach simply set all invertebrate taxa detected in the field to have equal abundance, to provide a baseline against which to assess the effects of different prey abundance estimates. When generating the two combined data sets, it was apparent that simply adding them together would underrepresent one of the datasets as abundance and activity density are measured in different units. Therefore, a ‘proportional combined’ dataset was generated by converting counts to relative proportions of each sample (to equally weight the two methods), which were then combined by summing proportions between the two methods for each sample, multiplied by the total count of individuals across both methods for each sample (to create realistic abundance values), and then rounded to the nearest integer (to return count data). In addition, a ‘frequency of occurrence (FOO) combined’ dataset was generated by converting counts to binary presence-absence values of each sample, which were then summed between the two methods for each sample. To assess the diversity represented by the two sampling methods and their combinations, and the completeness of those datasets, coverage-based rarefaction and extrapolation were carried out, and Hill diversity calculated (Chao et al., 2014; Roswell et al., 2021) using the ‘iNEXT’ package with families represented by frequency-of-occurrence across samples (Chao et al., 2014; Hsieh et al., 2016; Figures S1-S3).

**Figure 1:**
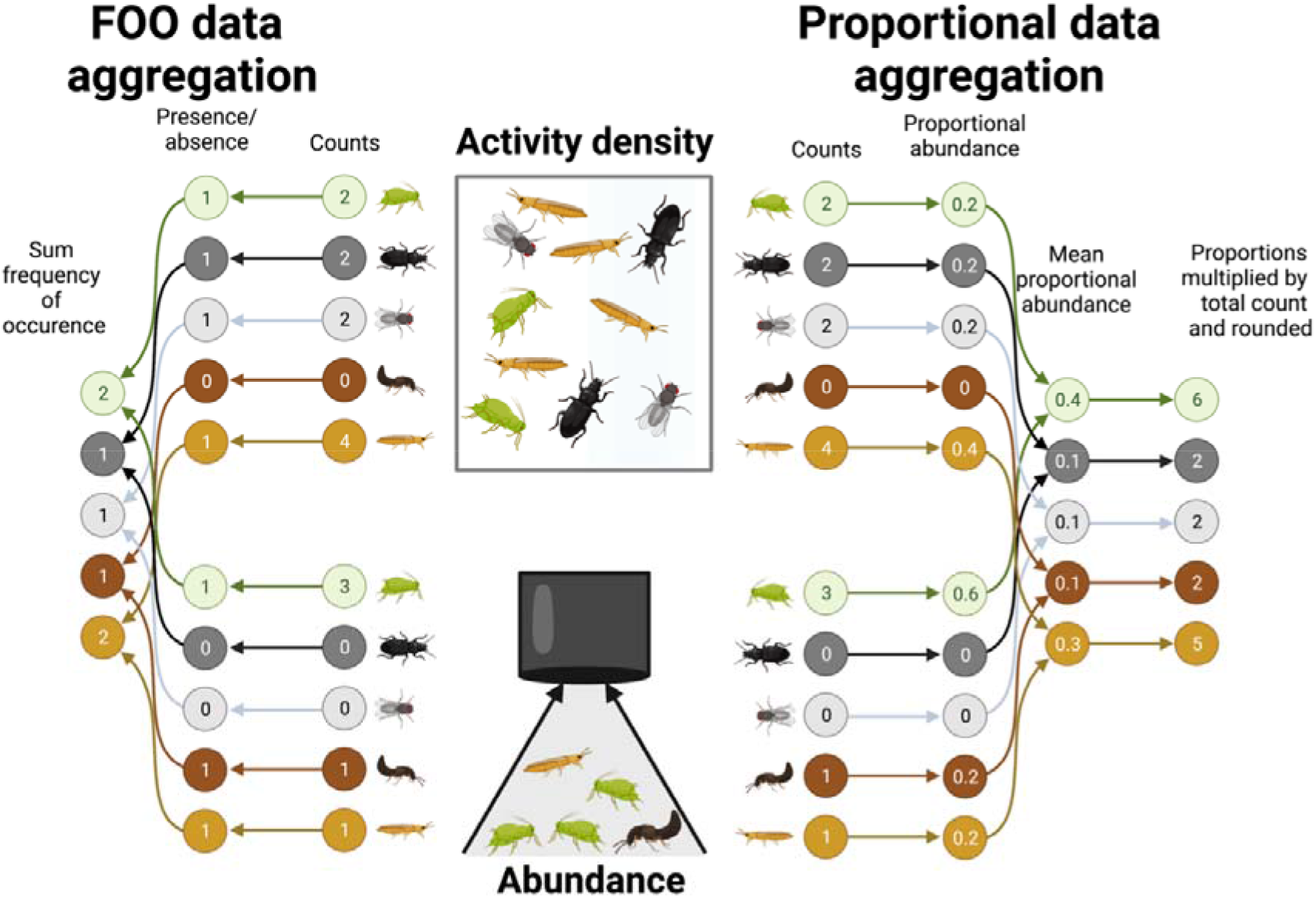
Data processing for the two combined method datasets from the original sticky trap and suction sample data. Figure created with Biorender.

The remaining analyses were performed using both presence-absence and relative read abundance dietary data separately to show how differences in the treatment of the observed data are reflected in the outcomes of the analyses. Figures and outputs given in the main text relate to the presence-absence data, while relative read abundance figures and outputs are presented in the Supplementary Information. Prey preferences of spiders were analysed using network-based null models in the ‘econullnetr’ package (Vaughan et al., 2018) with the ‘generate_null_net’ function. Econullnetr generates null models based on prey availability to predict how consumers would forage if based on the availability of resources alone. These null models are then compared against the observed interactions of consumers (e.g., interactions of spiders with their prey based on dietary metabarcoding) to ascertain the extent to which resource consumption deviated from random. In five separate null models, prey availability was represented separately by the datasets described above: abundance (suction sampling), activity density (sticky trapping), proportional combined, FOO combined and equal prey abundance.

To compare effect sizes between null models for each resource taxon, mean prey preference standardised effect size (SES) values were calculated from the individual spiders per model. The SES values were plotted and joined between taxa to visualise paired differences using ‘ggplot’ (Wickham, 2016). Null model-predicted trophic interactions were generated via an econullnetr null model with 999 simulations with outputs extended to allow the comparison of the null interactions for individual consumers (generate_null_net_indiv; Cuff, Kitson, et al., 2023). A visualisation of the per-individual differences in null model and observed data was generated via non-metric multi-dimensional scaling (NMDS) using the ‘metaMDS’ function in the ‘vegan’ package (Oksanen et al., 2016) in two dimensions and 9999 simulations, with Euclidean distance. Centroid coordinates for each null model and the observed data were extracted and pairwise distances calculated between model centroids:

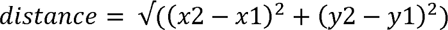

The ‘observed’ network (i.e., the network determined solely by dietary data, not necessarily the objectively ‘true’ network) and each null network were visualised with the associated prey choice effect sizes as a bipartite network using ‘ggnetwork’ (Briatte, 2021; Wickham, 2016) via an ‘igraph’ object (Csardi & Nepusz, 2006). The degree of each prey node, weighted nestedness and linkage density were generated using the ‘bipartite’ package (Dormann et al., 2008) for each network and compared visually via ggplot2.

## Results

### Dataset description

The dietary dataset used in this study contains data from 70 individual spiders which cumulatively interacted with 25 prey families, totalling 142 individual detected interactions. Sampling datasets contain data from 19 locations, with prey abundance data, determined via suction sampling, including 4766 individual invertebrates across 61 families (93.9 % complete; Figure S1) and prey activity density data, determined via sticky trapping, including 3513 individual invertebrates across 55 families (91.5 % complete; Figure S2). The prey availability data gained by combining the two measures of prey availability includes 85 families (95.1 % complete; Figure S3). The five prey availability datasets were relatively distinct and varied in their compositional similarity to the directly detected dietary data, with FOO combined being the most similar (Bray-Curtis distance = 0.545), followed by abundance (0.847), proportional combined (1.136), equal prey abundance (1.245) and activity density (1.362), respectively (Figure 2).

**Figure 2:**
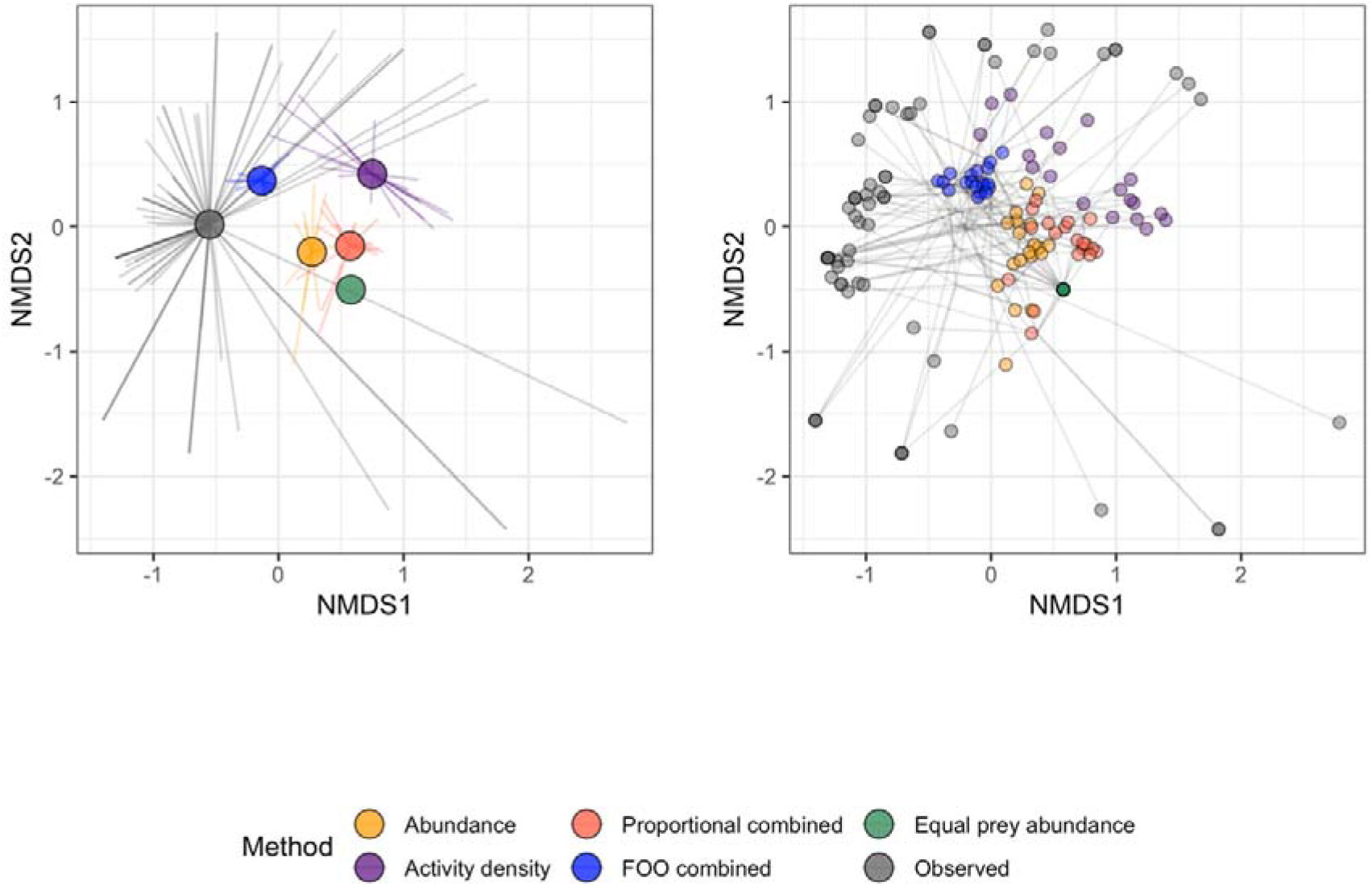
Spider plot derived from NMDS of spider diets (‘observed’) and prey communities derived from different sampling methods. In the left plot, the terminal end of each line represents a prey community or diet of a spider, joined by the centroids of diets from each data source (larger nodes; mean coordinates in that group), with distance between them indicating their dissimilarity (i.e., proximate points are similar, distant points are dissimilar). In the right plot, each point represents a prey community or diet of a spider, with lines joining points representing the same sample and those lines meeting at the mean coordinates for that sample.

### Differences in inferred foraging ecology

Predator selectivity differed substantially between datasets (Figure 3; Figure S6). The equal prey abundance, proportional combined and FOO combined datasets generated no significantly negative effect sizes (i.e., avoidances). The two combined datasets generated selectivity results largely consistent with the abundance data, but the FOO combined data generated null networks with the fewest significant deviations from the observed data. The activity density and abundance data showed some consistency, but sometimes generated opposite patterns and they tended to determine more significantly negative and positive deviations from observed data, respectively. The effect sizes inconsistently differed between datasets (Figure S4; Figure S7).

**Figure 3:**
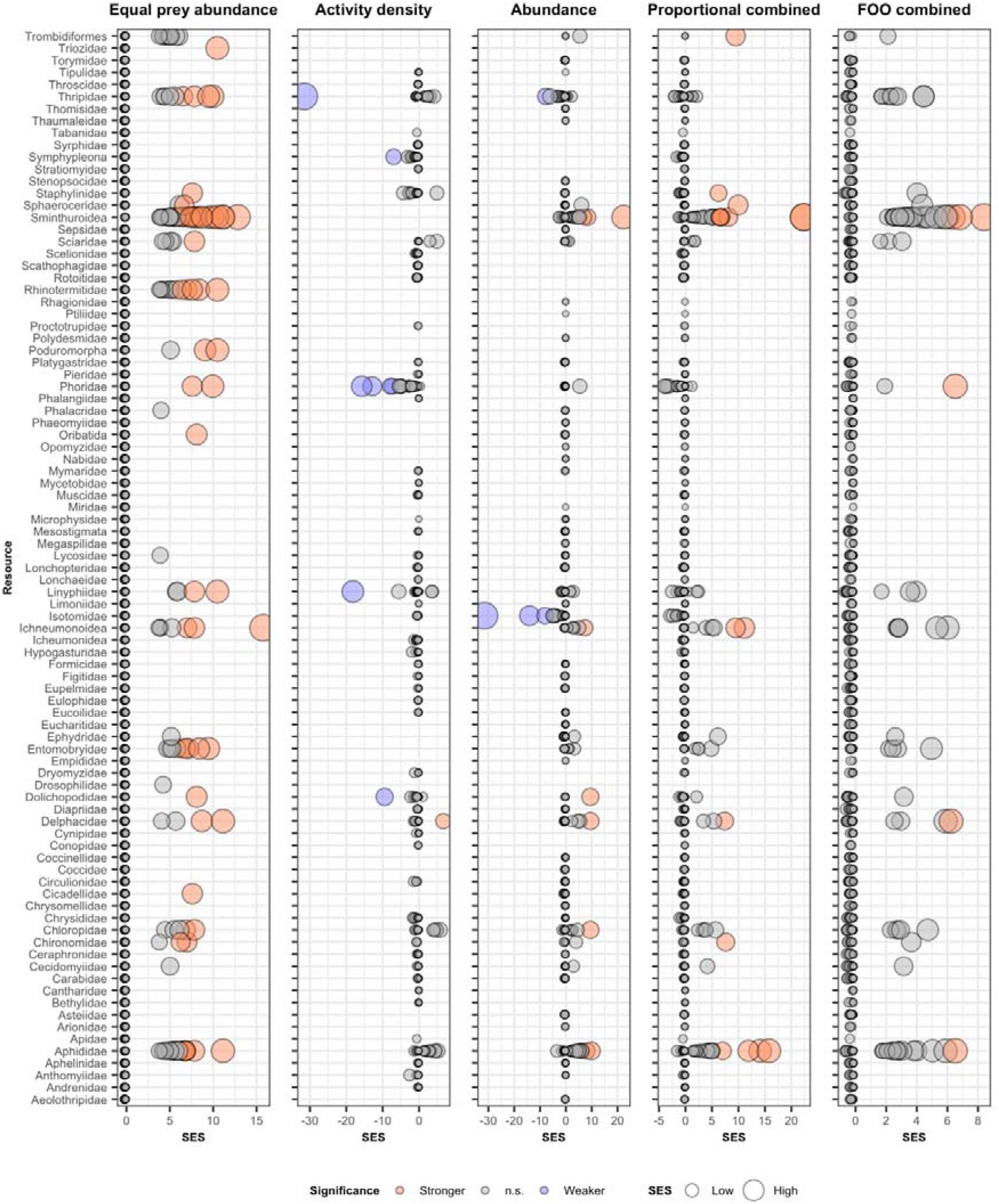
Prey choice standardised effect sizes (SESs) for each prey taxon and individual consumers for the five different null models. Larger points reflect larger deviations of SESs from zero (i.e., stronger or weaker selection). Red, white and blue points denote significantly more frequent (stronger selection), non-significant and significantly less frequent (weaker selection) interactions compared to the null model (*p* <0.05). Absent points are those for which data were not available.

### Differences in null network dietary compositions

The different prey availability datasets produced compositionally distinct null diets. The mean Euclidean coordinates of the diets generated via NMDS differed in their distance from the mean observed dietary composition (Figure 4; Figure S8), with FOO combined (Euclidean distance = 0.062), abundance (0.080) equal prey abundance (0.082), proportional combined (0.143) and activity density (0.269) being progressively further from the observed diets, respectively. All but abundance followed a relatively linear progression of difference from the observed diets with respect to NMDS axes.

**Figure 4:**
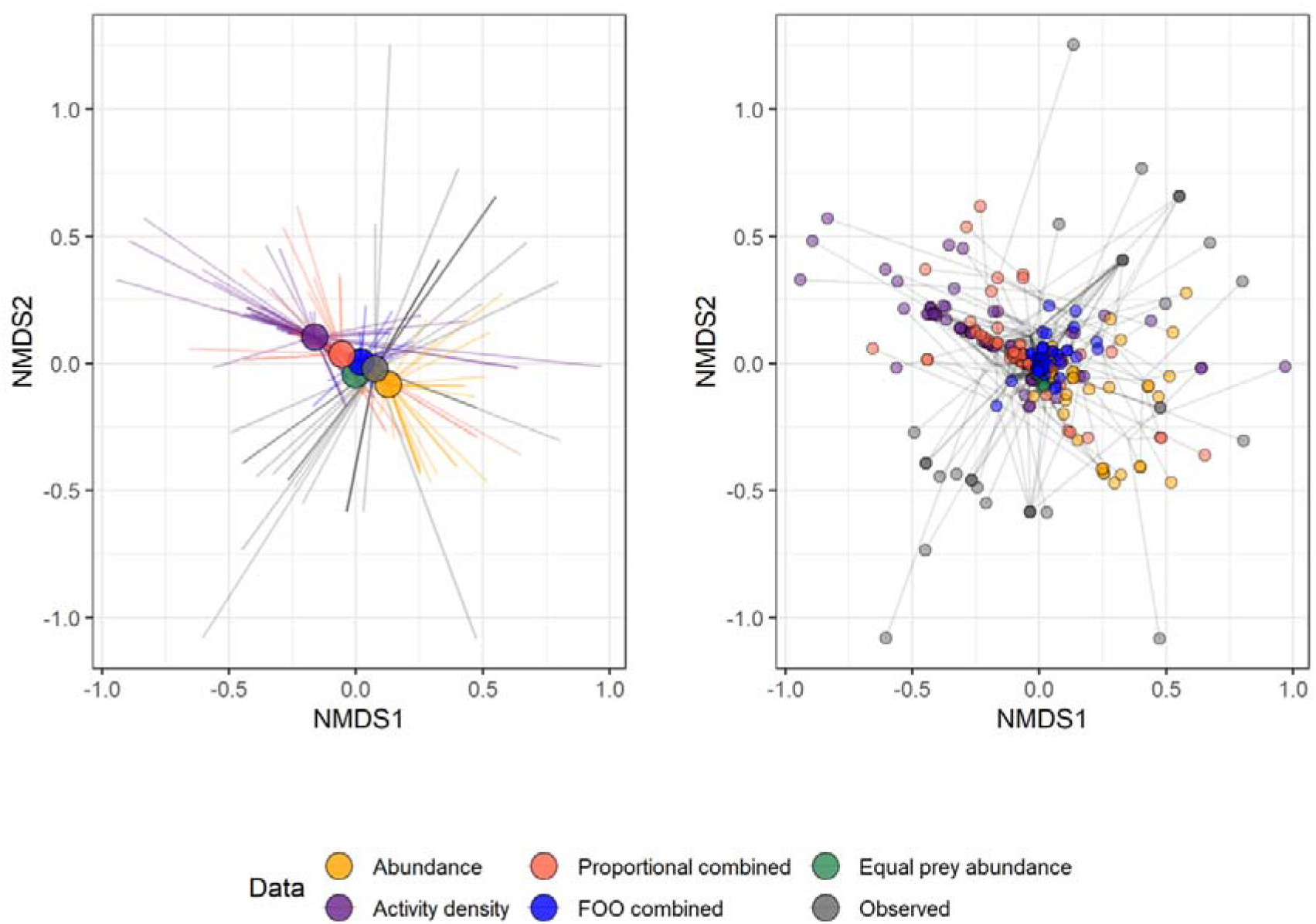
Spider plot derived from NMDS of observed and null model-expected spider diets. In the left plot, the terminal end of each line represents the diet of a spider (whether observed or simulated), joined by the centroids of diets from each data source (larger nodes; mean coordinates in that group), with distance between them indicating their dissimilarity (i.e., proximate points are similar, distant points are dissimilar). In the right plot, each point represents the diet of a spider, with lines joining points representing the same sample and those lines meeting at the mean coordinates for that sample.

### Differences in null network structure

The properties of null networks generated using the different prey availability datasets differed substantially (Figures 5-6; Figures S9-10). Nestedness and linkage density of the abundance, activity density and proportional combined null networks more closely resembled that of the network directly generated from dietary data (Figure 6; Figure S10), as did the degree of prey nodes in many instances (Figure S5; Figure S11).

**Figure 5:**
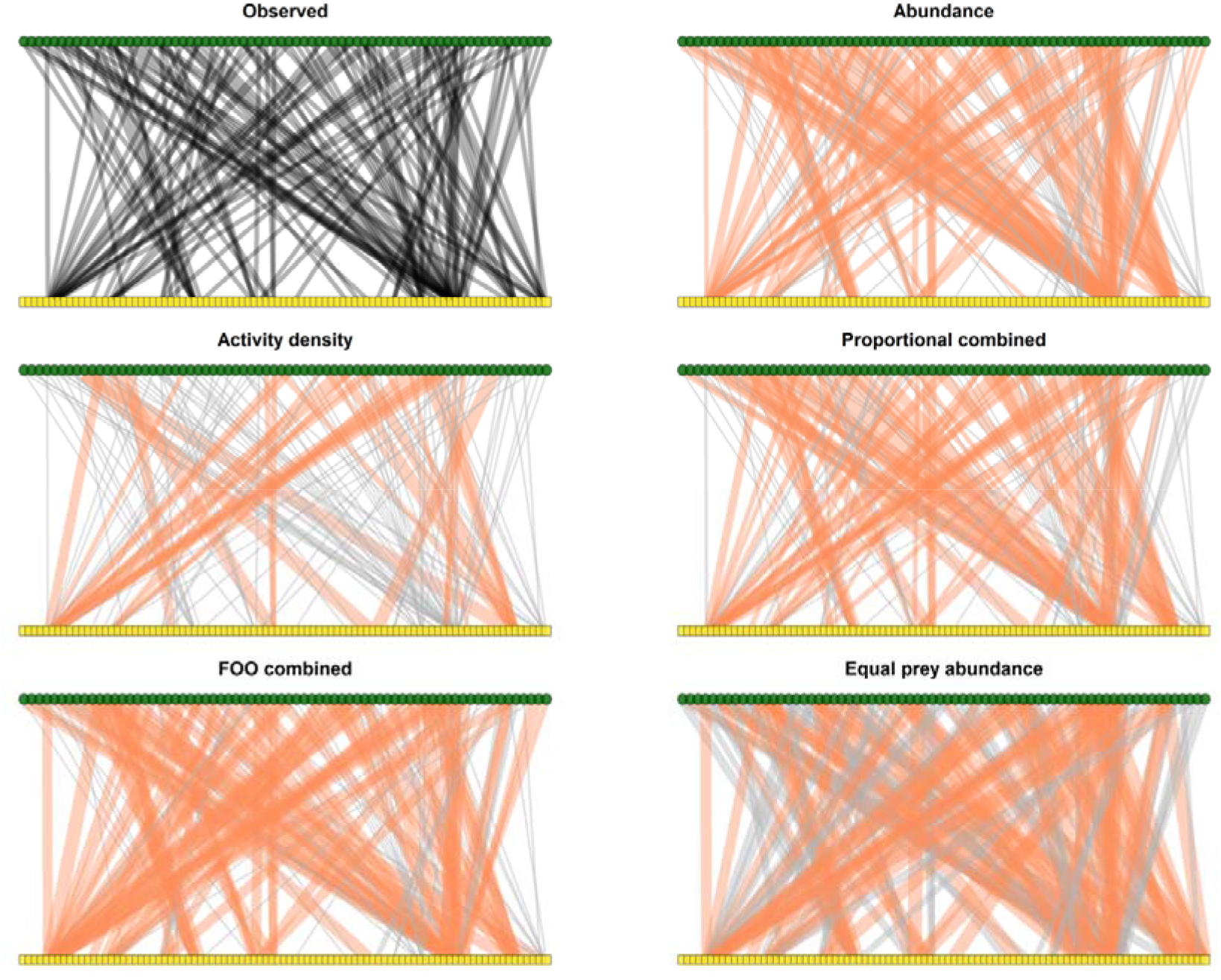
Network generated solely from dietary data (‘observed’) and null networks produced from the different prey availability datasets. Link weights represent the number of observed interactions for the ‘observed’ network, and otherwise the frequency of interactions expected based on prey availability according to null models. Red links represent those for which observed interaction frequencies significantly exceeded those expected from null models. No interactions are plotted which were significantly less frequent than expected.

**Figure 6:**
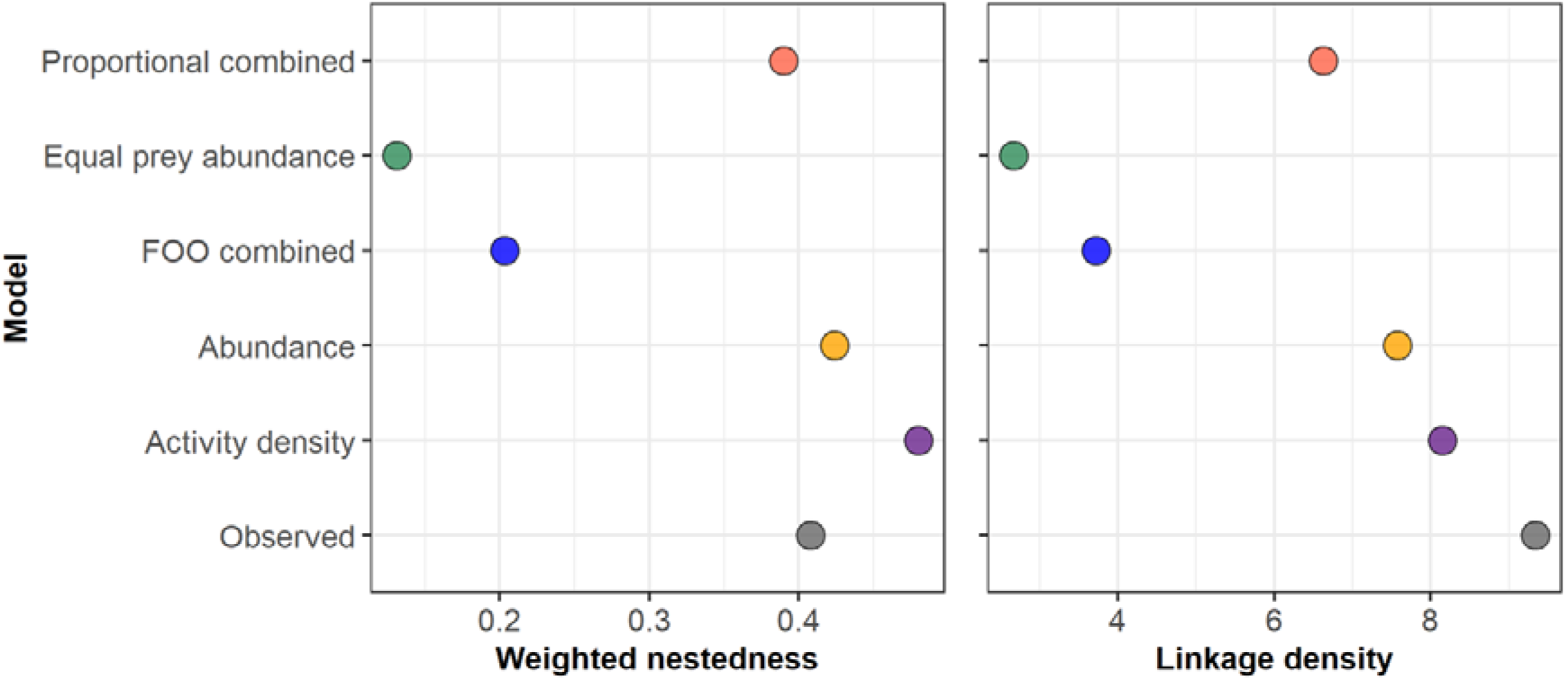
Weighted nestedness and linkage density of the ‘observed’ network and the null networks generated from the different prey availability datasets.

## Discussion

Prey sampling method has implications for the interpretation of predator selectivity, dietary composition and network structure when using null networks, but the precise nature of these effects is complex. The implications of prey sampling method are dependent on the specific questions being addressed, with the methods for estimating prey availability having different effects upon the frequency of interactions, dietary composition and network structures of null networks and, consequently, the inferred selectivity of predators. Increasing the completeness of datasets by merging sampling methods could theoretically better simulate trophic interactions by increasing the accuracy of prey availability data. We show, however, that combining methods is not straightforward and may generate null diet compositions or network structures that differ more from those generated from directly detected interactions, depending on the method of merging. Even though relying on single biased methods can misrepresent true prey availability, we lack methods for reliably improving these estimates. Because all methods for estimating prey availability have their biases, we must always treat estimated predator selectivity and corresponding network metrics with caution. The optimal representation of prey availability in these contexts depends on the hypotheses being tested and, most crucially, the ecology of the system being studied.

### Considerations when choosing sampling methods for null network choice analysis

We have shown that different sampling methods affect the outcomes of predator selectivity analyses. This conclusion is intuitive since altering the data from which null models are generated will naturally change the structure of the null networks and the identities of the resource nodes within. The effects are, however, far more nuanced and differ depending on the characteristics being assessed. Some sampling methods generated null networks with structural properties more like the network constructed using directly detected interactions, and other sampling methods generated null diets with prey identities and frequencies more like the directly detected network. This finding predicates that optimal sampling method choice is not only system-dependent, but also contingent upon the specific hypotheses being tested.

Sampling methods have well-documented and characterised taxonomic and functional biases which can vary greatly, as exemplified by the methods used in this study. Whilst suction sampling can capture many of the smaller near-ground prey commonly exploited by spiders (Cooper & Whitmore, 1990; Harper & Guynn, 1998), it can disproportionately represent thrips, spiders, true bugs, flies and wasps (Doxon et al., 2011; Zentane et al., 2016). The ‘peripheral suction effect’, whereby some taxa are predisposed to being collected from beyond the sampling area, exacerbates these taxonomic biases, although it can be mitigated by surrounding the sampling area with a cylinder (Cherrill, 2015). Other factors, such as time of day, weather conditions and time spent suction sampling may also influence the outcomes (Bell et al., 2000; Brook et al., 2008). Similarly, sticky traps elicit different biases based on the colour used, which determines attraction of different taxa (Böckmann & Meyhöfer, 2017; Chittka & Menzel, 1992; Döring et al., 2012; Hoback et al., 1999; Sétamou et al., 2014). Ultimately, all sampling methods are biased, but when representing the availability of prey to a predator, the sampling method should ideally emulate, as closely as possible, the foraging behaviour and prey capture opportunities of the predator (Table 1).

In a hypothetical study concerning the prey preferences of trapdoor spiders (Araneae: Halonoproctidae), the researchers involved should consider the foraging mode of the predator to select an appropriate prey survey method. Given that trapdoor spiders are sit-and-wait predators, it is likely that activity density would better reflect the prey available to the spiders. Since the trapdoors of these spiders, from which they ambush prey, resemble pits in the ground, pitfall traps may be an intuitive choice of sampling method. In fact, one study highlights that pitfall trap abundances correlate with trapdoor spider activity (Bradley, 1996), suggesting that this would be an appropriate choice. Through reference to existing literature and natural history records (e.g., Bradley, 1996; Coyle & Icenogle, 1994; Gupta et al., 2015), the researchers could identify the likely prey of trapdoor spiders and whether this method will collect them. Further considerations like the diameter of the trap opening (Table 1) could also be adjusted according to existing data within the system to design the most realistic representation of prey capture. In the case of the trapdoor spider, prey vibrational cues instigate predation (Nakamura et al., 2022), thus the prey collected during surveys must also be considered from this physiological perspective (i.e., whether they would produce a viable vibrational cue to trigger their predation) unless this is the mechanism of choice being explored via null modelling.

Datasets generated by combining data from both sampling methods were more complete and included a broader spectrum of available prey, leading to fewer false negatives in null models (i.e., prey that were detected in the guts of spiders, but not found in the prey availability data). Due to the greater imbalance in prey counts in these datasets, however, interaction frequencies were distributed across a greater number of prey, leading to less realistic network topologies and interaction weights. These ‘false positive’ null interactions which did not occur in the directly detected interaction data could be mitigated by restricting resource availability data to only those taxa with which consumers were found to interact, but this would obviate any investigation of the mechanisms potentially driving the exclusion of these taxa from consumer interactions. Merging sampling data from different survey methods is thus a complicated solution that may induce additional biases; whilst it overcomes individual methodological biases and increases sampling completeness, it may inflate deviation of directly detected network properties from null models. This could equally be due to an unaccounted for ecological phenomenon though. Inundating null models with available prey will nevertheless distribute interactions across a broader range of resources, potentially reducing linkage density, modularity and nestedness. Poorly informed data-saturated null networks in which any consumer-resource interaction is permitted will thus poorly represent a baseline against which to compare real-world trophic interactions in which consumers are time, energy and resource limited and will thus be more selective. Care must be taken to ensure that only plausible interactions are represented by the resultant null models, or those pertaining to the hypothesis being tested.

It is important to consider how survey data align with the observed data with which they are being compared. Directly detected or observed network data are often subject to biases and limitations with similar implications for differences between observed and null networks. For example, the time scales over which observations and prey survey data are collected may mismatch if the prey with which predators interact are no longer present at the time of surveying (e.g., the long period of detection of DNA data compared to active prey abundance surveys). This problem is particularly noteworthy for prey present at different times in the diel cycle which may be underrepresented by abundance data. It is thus crucial to consider how measures of observed interactions and prey availability data align during experimental design.

Dietary metabarcoding data, increasingly used in null models assessing resource choice (Cuff et al., 2021; Cuff, Tercel, Drake, Vaughan, et al., 2022; Davies et al., 2022; Evens et al., 2020; Gajski et al., 2023; Moorhouse□Gann et al., 2022; Verschut et al., 2019; Villsen et al., 2022), also present several distinct considerations. Alongside lacking the context of prey life stage, sex and other contextual information that such null models could otherwise include to assess how prey traits affect foraging, quantification of metabarcoding data is a vital concern. Trophic interactions represented as binary presence-absence data lack realistic interaction weights, whereas relative read abundances generated by metabarcoding present quantities, but they are often taxonomically biased (Deagle et al., 2019; Lamb et al., 2019). Neglecting quantitative data will, at least at the individual level, mismatch the weighted interactions of null models, whereas introducing biased data could poorly reflect realistic foraging. By running these analyses with both data types (Figures S6-S11), we have shown that the overall differences between observed data and corresponding null networks were relatively consistent regardless of the observed data used. The key differences were that relative read abundances generated more significant deviations between null and observed data, but that these were largely just stronger effect sizes for the same taxa highlighted by presence-absence-based analyses. This demonstrates that the core findings apply regardless of the observed data used.

Dietary metabarcoding data are also subject to false positives (Drake et al., 2022) and false negatives (Littleford□Colquhoun et al., 2022). This is particularly insidious for omnivorous consumers (Tercel et al., 2021) but can be exacerbated by amplification of the DNA of the focal predator itself (Cuff, Kitson, et al., 2023). The presence of false positives/negatives undoubtedly alters the congruence of observed data with null networks and the inferred strength of resource preferences, thus a careful approach that limits data loss whilst preserving data integrity is required (Littleford□Colquhoun et al., 2022). By merging networks constructed with DNA and direct observation data (i.e., visually recorded interaction data), some of the biases and limitations of both methods can be overcome (Cuff, Windsor, Tercel, Kitson, et al., 2022), but how this affects congruence of dietary and prey availability data may vary greatly. Ultimately, the methodological biases imposed on the prey availability and observed interaction data should align as much as possible to limit the impact of dataset mismatches on ecological outcomes.

### Example of the ecological information gained from null model comparisons

To assess the relevance of different measures of prey availability, it is important to consider what these measures should achieve. The aim of null modelling is to investigate specific mechanisms as drivers of ecological patterns through comparison of observed data with data generated according to a specific null hypothesis (Gotelli, 2001; Gotelli & Graves, 1996). In the context of prey choice, the models used in this study are purposed to identify interactions that occur more or less frequently than would be expected if predators randomly sampled from the community of prey available to them (Vaughan et al., 2018). The objective of sampling is therefore to represent the prey available to each predator, for which the ecology of each forager is a vital consideration.

The predators in this study have variable foraging behaviour and ecology, best represented by the two Linyphiidae subfamilies Linyphiinae and Erigoninae. These spiders use webs to forage, the position and size of which differs even within families (Harwood et al., 2001). Linyphiinae spiders build larger sheet webs a few centimetres above the ground, whereas Erigoninae spiders build smaller webs closer to the ground which they leave regularly to forage (Sunderland et al., 1986), lending to separation of their trophic niches (Harwood et al., 2003). Given the difference in active and passive foraging, it might be predicted that the interactions of these spider groups differed in their similarity to null networks generated using abundance and activity density data, with Erigoninae resembling abundance, and Linyphiinae activity density.

The interactions of all spider groups more closely resembled abundance than activity density, and the interactions of Erigoninae, whilst more like the abundance-based null network, were much more similar to the activity density null diets than Linyphiinae (Figure S12). This difference may indicate that neither method, nor their combinations, perfectly reflected the availability of prey to Linyphiinae. Other methods were trialled, including longer ground-pinned sticky traps, and acetate sheets coated with ecological glue (Oecotak adhesive) which approximately matched the webs of each spider in size and position, both of which often captured too few intact prey to generate null networks. Alternatively, the low congruence of Linyphiinae interactions with null networks may suggest that Linyphiinae simply have stronger density-independent preferences than Erigoninae (see Cuff, Tercel, et al., 2022 for an in-depth taxonomic comparison of preferences), as these methods are intended to determine. The latter case would highlight that, whilst null networks need to accurately represent prey availability, similarity between null networks and networks derived from directly detected or observed interactions does not absolutely equate to the suitability of the survey method given that density-independent foraging is commonplace. Perfectly simulating the diet of the consumer, whilst useful for predictive applications (Cuff, Windsor, et al., 2023), is not usually the aim of null networks used in this context. Instead, null networks should represent interactions that are physiologically, spatially and temporally accessible to the consumer. As such, observed interactions can inform which sampling method to deploy in specific contexts, but they cannot govern these decisions without introducing dogma.

The equal prey abundance null diets were compositionally more similar to the observed diet than those based on activity density or proportionally combined data, indicating that spiders achieved relatively even interactions across the diversity of prey available, arguably irrespective of prey activity density. This pattern suggests that prey community diversity is a greater driver of interaction diversity than prey activity densities for these spiders, which has important implications not only for optimal foraging theory, since it suggests foraging within optimal patches rather than for optimal prey (MacArthur & Pianka, 1966), but also for the ‘iDNA’ (invertebrate-derived DNA) monitoring concept (Cutajar & Rowley, 2020; Drinkwater et al., 2021) since it highlights the potential validity of using invertebrates as samplers of DNA for diversity assessment.

## Conclusions

We have demonstrated the substantial influence of different measures of prey availability on ecological interpretation of null network-based prey choice analyses. We also have shown that the choice of survey method should be dependent on the hypotheses, study system and predator foraging mode being investigated. Merging different data types increases data completeness and can produce null networks similar to those constructed from directly detected interactions, but the networks, interactions and null diets generated were similar to those generated with data from a single measure of prey abundance. During experimental design, researchers must carefully consider the foraging mode and ecology of the focal predator to best represent their available prey, but also the breadth of prey that they are likely to interact with and the concomitant network structures. Increasingly data-rich approaches to null network analysis have further implications for methodological choices. Integrating data such as environmental context (Cuff, Windsor, et al., 2023) and consumer functional trait data (Ibanez, 2012) can refine or test predictions regarding consumer choices and could greatly expand the utility and benefit of null modelling approaches, but robust resource availability data must be sought in all such applications.

## Acknowledgements

Thanks to Robert Reader for use of Burdons Farm, Wenvoe for fieldwork. JPC was funded by the Biotechnology and Biological Sciences Research Council through the South West Biosciences Doctoral Training Partnership (grant BB/M009122/1). JRB was funded by BBSRC’s Core Capability Grant (BBS/E/C/000J0200). MPTGT was funded by the Natural Environment Research Council (NE/L002434/1) and Durrell Wildlife Conservation Trust (MR/S502455/1).

## Data availability

All data and R scripts are openly available at Zenodo: https://doi.org/10.5281/zenodo.7908186 (Cuff, Tercel, et al., 2023)

## Author contributions

JPC, MPTGT and IPV conceived the core ideas and oversaw the project with additional ideas and conceptual contributions from BSJH and PAH; JPC, MPTGT, FMW and IPV analysed the data; WOCS, IPV, JRB and JPC acquired funding; JPC led writing of the manuscript with substantial contributions from all authors.

