## Supplementary Information for "Sources of prey availability data alter interpretation of outputs from prey choice null networks"

### Prey availability data sources alter ecological interpretation from null network prey choice analyses – supplementary materials

**Supplementary Information 1: Molecular analysis and bioinformatics**

##### *Extraction and high-throughput sequencing of spider gut DNA*

Given their prevalence in field collections, dietary analysis was carried out for the linyphiid genera *Erigone*, *Tenuiphantes*, *Bathyphantes* and *Microlinyphia* (Araneae: Linyphiidae), and the Lycosidae genus *Pardosa*. Spiders were transferred to and washed in fresh 100 % ethanol to reduce external contaminants prior to identification via morphological key (Roberts, 1993). Abdomens were removed from spiders and again transferred to and washed in fresh 100 % ethanol. DNA was extracted from the abdomens via Qiagen TissueLyser II and DNeasy Blood & Tissue Kit (Qiagen) as per the manufacturer protocol, but with an extended lysis time of 12 hours to account for the complex and branched gut system in spider abdomens (Krehenwinkel et al., 2017).

For amplification of DNA, two primer pairs were used. BerenF-LuthienR (Cuff et al., 2021) amplified a broad range of invertebrates including spiders, and TelperionF-LaureR (Cuff et al., 2022), amplified a range of invertebrates but fewer spiders. Primers were labelled with unique 10 bp molecular identifier tags (MID-tags) so that each individual had a unique pairing of forward and reverse tags for identification of each spider post-sequencing. PCR reactions of 25 µl contained 12.5 µl Qiagen PCR Multiplex kit, 0.2 µmol (2.5 µl of 2 µM) of each primer and 5 µl template DNA. Reactions were carried out in the same thermocycler, optimised via temperature gradient, with an initial 15 minutes at 95 °C, 35 cycles of 95 °C for 30 seconds, the primer-specific annealing temperature for 90 seconds and 72 °C for 90 seconds, respectively, followed by a final extension at 72 °C for 10 minutes. BerenF-LuthienR and TelperionF-LaureR used annealing temperatures of 52 °C and 42 °C, respectively.

Within each PCR 96-well plate, 12 negative controls (extraction and PCR), 2 blank controls and 2 positive controls were included (i.e. 80 samples per plate), based on Taberlet *et al.* (2018). Positive controls were mixtures of invertebrate DNA comprised of non-native Asiatic species in four different proportions and blanks were empty wells within each plate to identify tag-jumping into unused MID-tag combinations. PCR negative controls were DNase-free water treated identically to DNA samples. A negative control was present for each MID-tag to identify any contamination of primers. All PCR products were visualised in a 2 % agarose gel with SYBRSafe (Thermo Fisher Scientific, Paisley, UK) and placed in categories based on their relative brightness. The concentration of these brightness categories was quantified via Qubit dsDNA High-sensitivity Assay Kits (Thermo Fisher Scientific, Waltham, MA, USA) with at least three representatives of each category per plate. The PCR products were then proportionally pooled according to these concentrations. Each pool was cleaned via SPRIselect beads (Beckman Coulter, Brea, USA), with a left-side size selection using a 1:1 ratio (retaining ~300-1000 bp fragments). The concentration of the pooled DNA was then determined via Qubit dsDNA High-sensitivity Assay Kits and pooled together into one library per primer pair. Library preparation for Illumina sequencing was carried out on the cleaned libraries via NEXTflex Rapid DNA-Seq Kit (Bioo Scientific, Austin, USA) and samples were sequenced on an Illumina MiSeq via a V3 chip with 300-bp paired-end reads (expected capacity ≤25,000,000 reads). Bioinformatic analysis followed Drake et al. (2022).

##### *Bioinformatic analysis*

The Illumina run generated 11,165,405 and 10,959,010 reads for BerenF-LuthienR and TelperionF-LaureR, respectively, which were quality-checked and paired via FastP (Chen et al., 2018) to retain only sequences of at least 200 bp with a quality threshold of 33, resulting in 10,561,874 and 9,355,112 paired reads. The paired reads were demultiplexed and assigned to their respective spider sample according to their MID-tags via the “trim.seqs” command in Mothur v1.39.5 (Schloss et al., 2009), leaving 7,854,610 and 7,437,929 reads with exact matches to the primer and MID-tags.

Replicates were removed, and denoising and clustering to zero-radius operational taxonomic units (ZOTUs; clustered without % identity to avoid multiple species represented within a single operational taxonomic unit (OTU)) completed via Unoise3 in Usearch11 (Edgar, 2010). The resultant sequences were assigned a taxonomic identity from GenBank via BLASTn v2.7.1 (Camacho et al., 2009) using a 97 % identity threshold (Alberdi et al., 2017). The BLAST output was analysed in MEGAN v6.15.2 (Huson et al., 2016). Where the top BLAST hit, determined by lowest e-value, was resolved at a higher taxonomic level than species-level, the results were checked; where possibly erroneous entries were preventing species-level assignment (e.g., poorly resolved identifications on GenBank), finer resolution was assigned based on the next-closest match. Where ZOTUs were assigned the same taxon, these were aggregated.

Data clean-up used the optimal minimum sequence copy thresholds identified by Drake et al. (2022). The maximum value for a ZOTU present in blank or negative controls was identified and subtracted from all read counts for that ZOTU to remove background contaminants. Simultaneously, known lab contaminants (e.g., German cockroach *Blattella germanica*), artefacts and errors of the sequencing process, unexpected reads in positive controls and positive control taxon reads in dietary samples were identified. These were calculated as a percentage of their respective sample’s read count and any read counts lower than the highest of these percentages for their respective sample were removed to eliminate additional instances of contamination. These thresholds were defined as 0.38 % and 0.39 % for BerenF-LuthienR and TelperionF-LaureR, respectively. The data from the two libraries (i.e., from each primer pair) were then aggregated together by sample and aggregated again by taxon. Non-target taxa (e.g., fungi) and instances in which predator DNA was amplified (i.e., ZOTUs with high read counts matching the individual’s morphological identity) were removed.

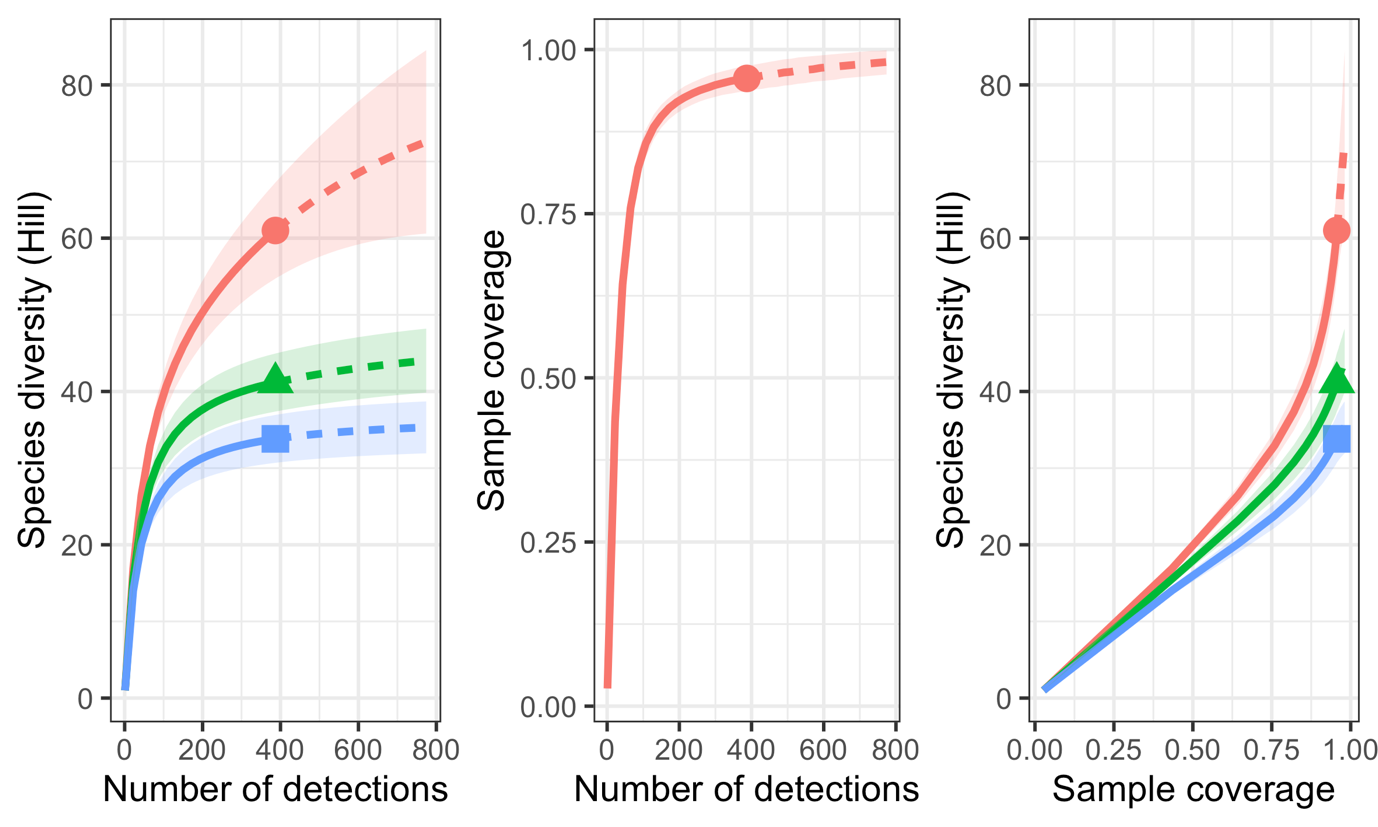

Figure S1: Diversity detected during suction sampling, and the associated sample coverage. Red lines with terminal circles, green with terminal triangles and blue with terminal squares denote Hill-species richness, Hill-Shannon and Hill-Simpson, respectively. Solid lines represent observed diversity, and dashed lines extrapolated diversity. Light zones surrounding lines denote 95 % confidence intervals.

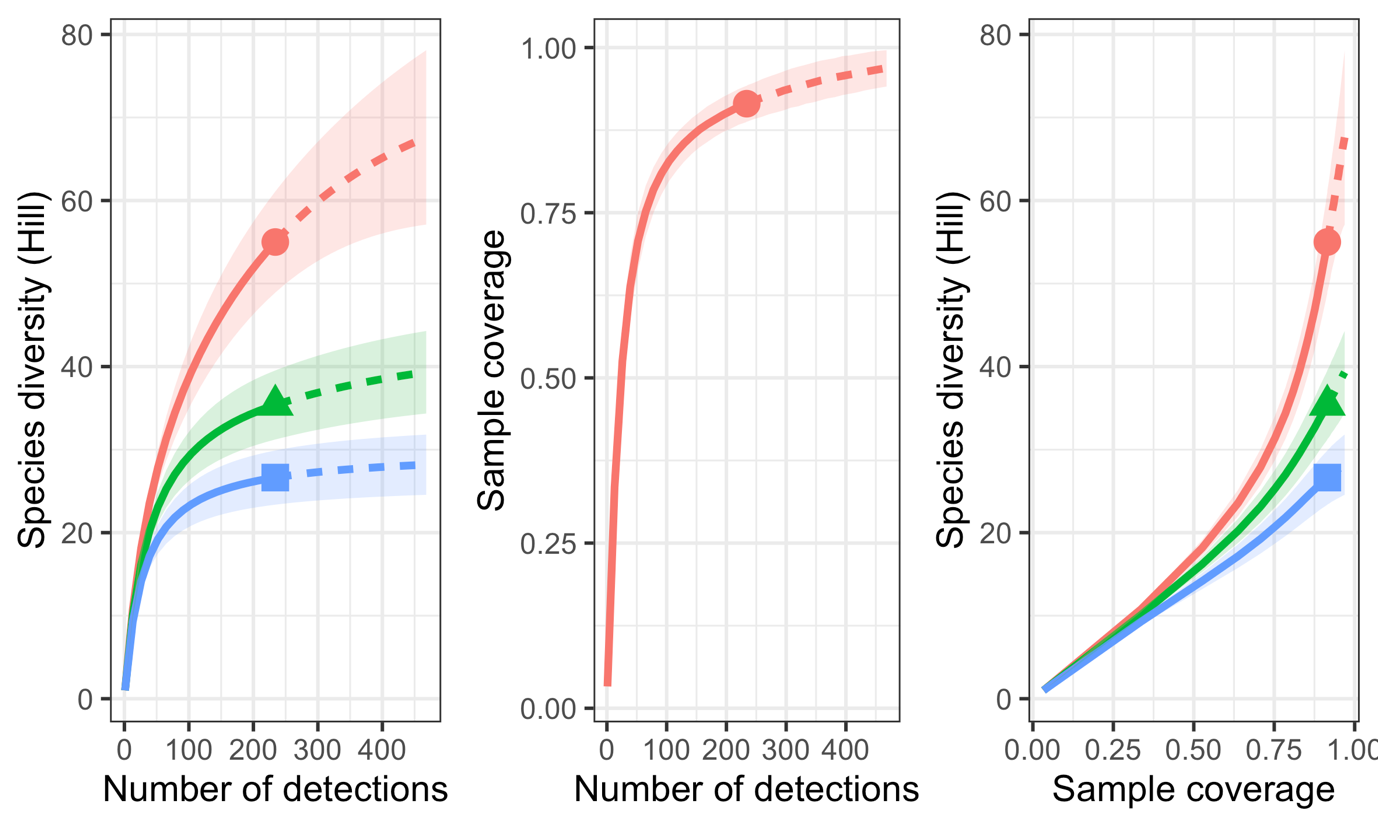

Figure S2: Diversity detected during sticky trapping, and the associated sample coverage. Red lines with terminal circles, green with terminal triangles and blue with terminal squares denote Hill-species richness, Hill-Shannon and Hill-Simpson, respectively. Solid lines represent observed diversity, and dashed lines extrapolated diversity. Light zones surrounding lines denote 95 % confidence intervals.

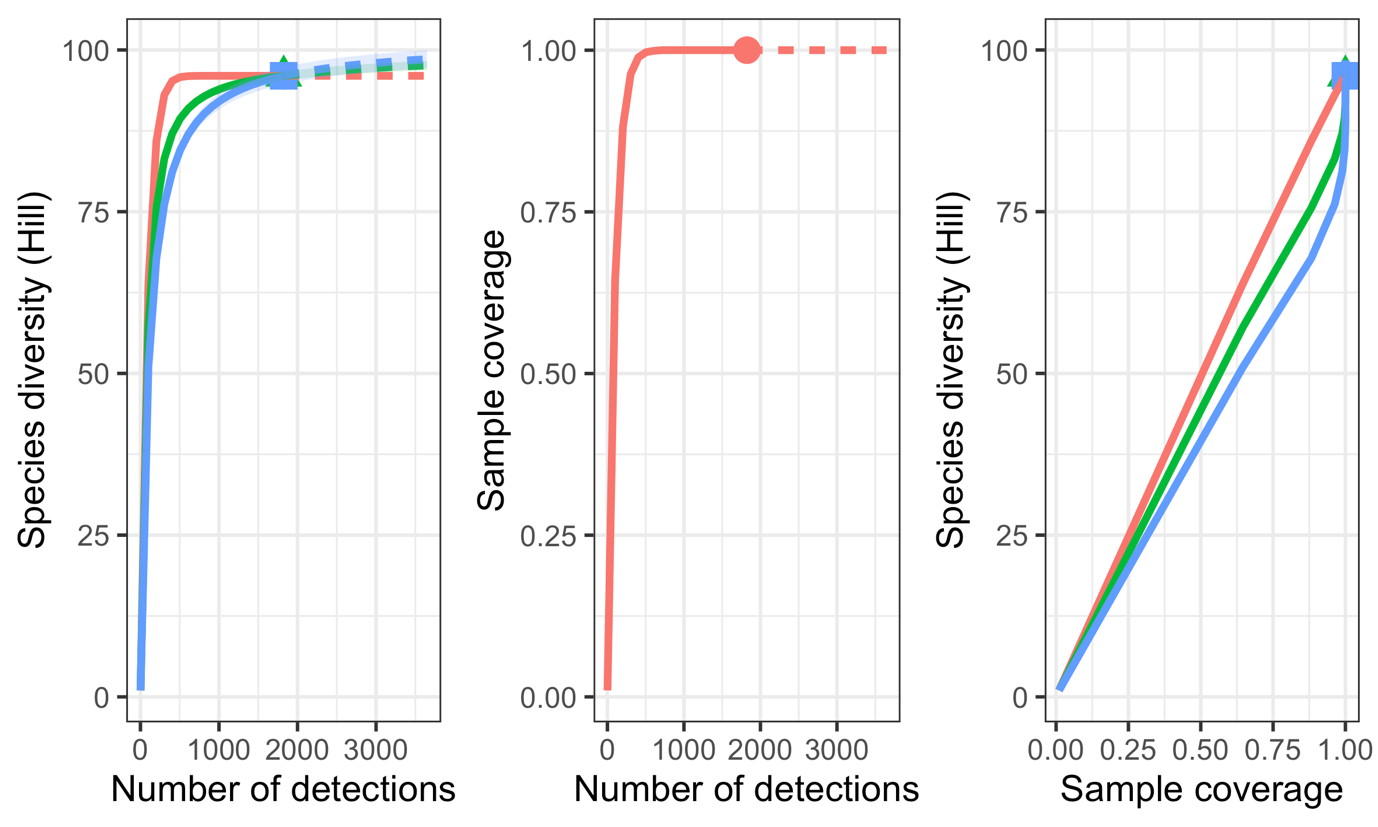

Figure S3: Diversity detected by the combination of suction sampling and sticky trapping, and the associated sample coverage. Red lines with terminal circles, green with terminal triangles and blue with terminal squares denote Hill-species richness, Hill-Shannon and Hill-Simpson, respectively. Solid lines represent observed diversity, and dashed lines extrapolated diversity. Light zones surrounding lines denote 95 % confidence intervals.

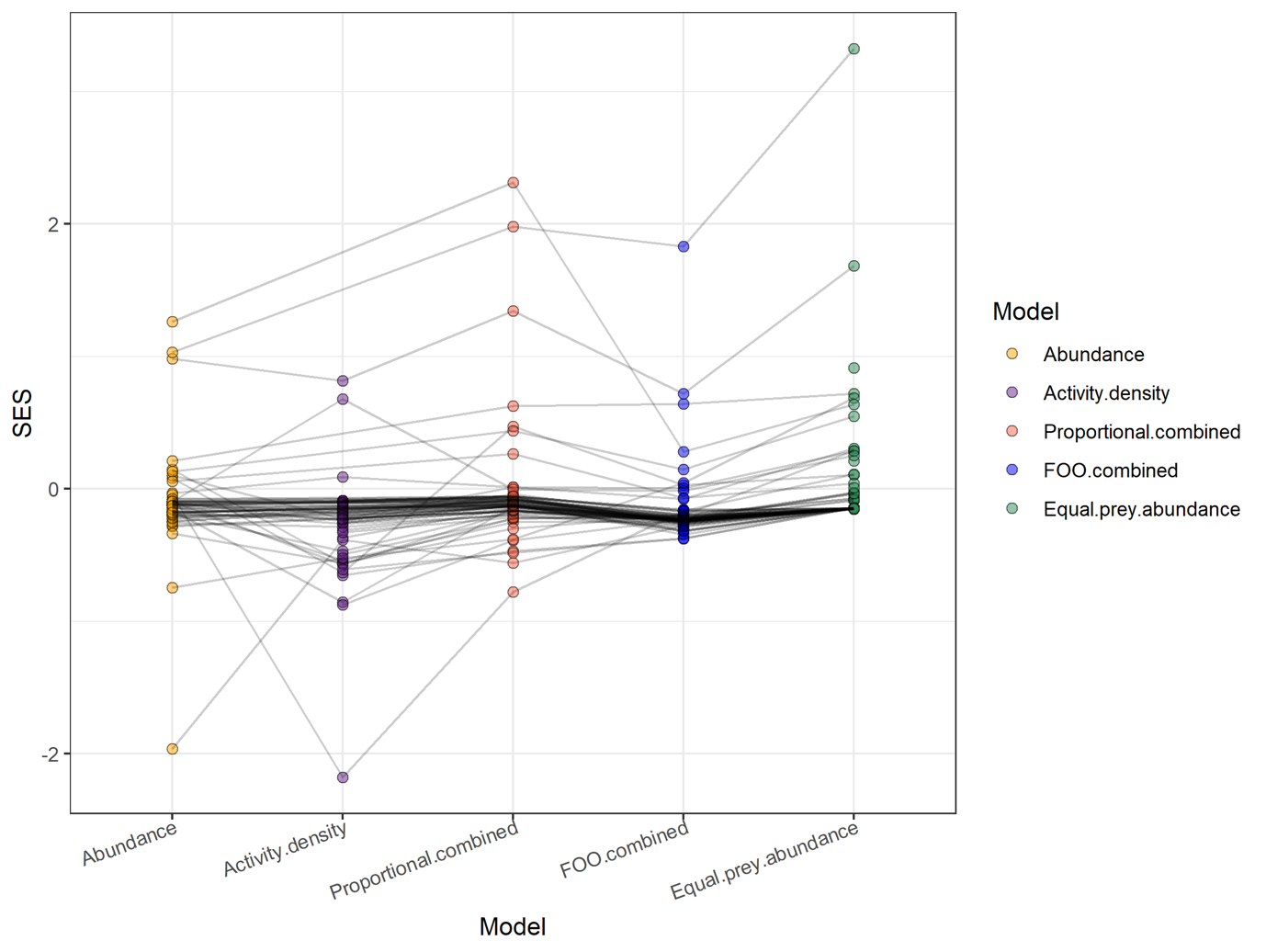

Figure S4: Standardised effect sizes from the comparison of null models against observed interactions. Mean SES values for each prey taxon across the individual consumers were used. Each colour denotes SESs from a different prey choice model. Links are drawn between points representing the same prey taxon.

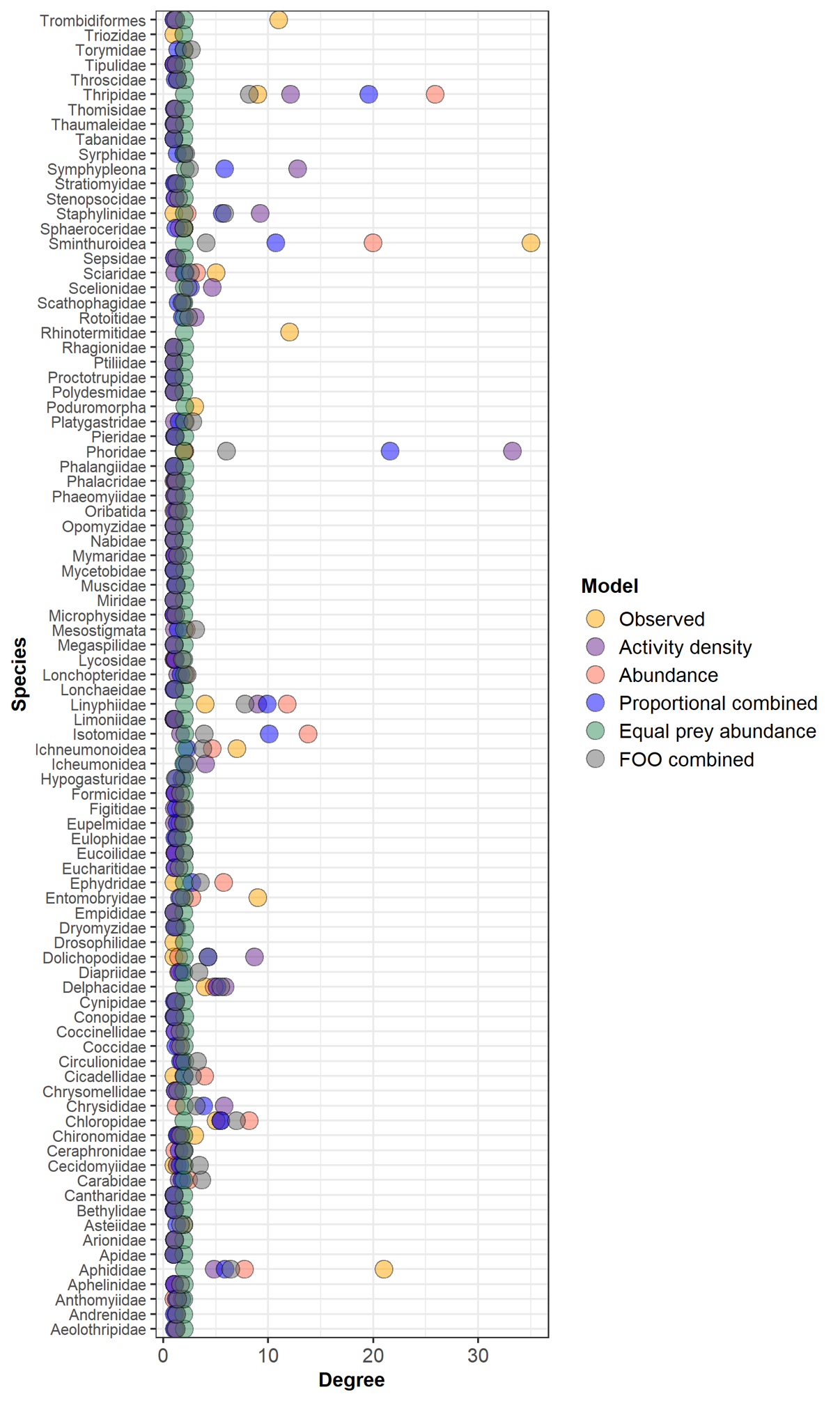

Figure S5: Degree of each node within the observed (presence-absence) and the null networks generated from the different prey availability datasets.

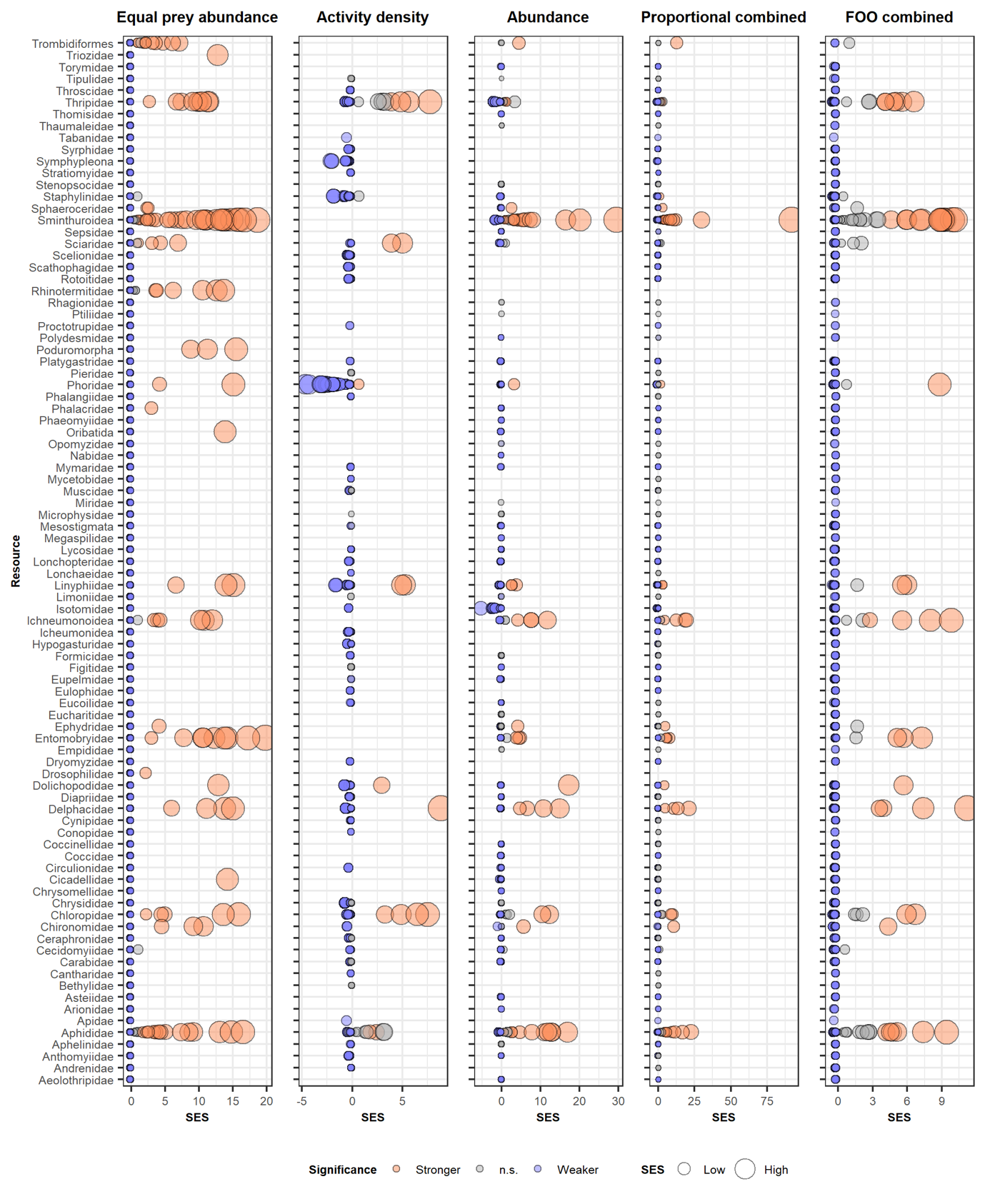

Figure S6: Relative read abundance-based prey choice standardised effect sizes (SESs) for each prey taxon and individual consumers for the five different null models. Larger points reflect larger deviations of SESs from zero (i.e., stronger or weaker selection). Red, white and blue points denote significantly more frequent (stronger selection), non-significant and significantly less frequent (weaker selection) interactions compared to the null model. Absent points are those for which data were not available. The overall patterns were consistent between presence-absence and relative read abundance data, but the latter identifies more significant deviations from the null model and stronger SESs generally (Figure 3).

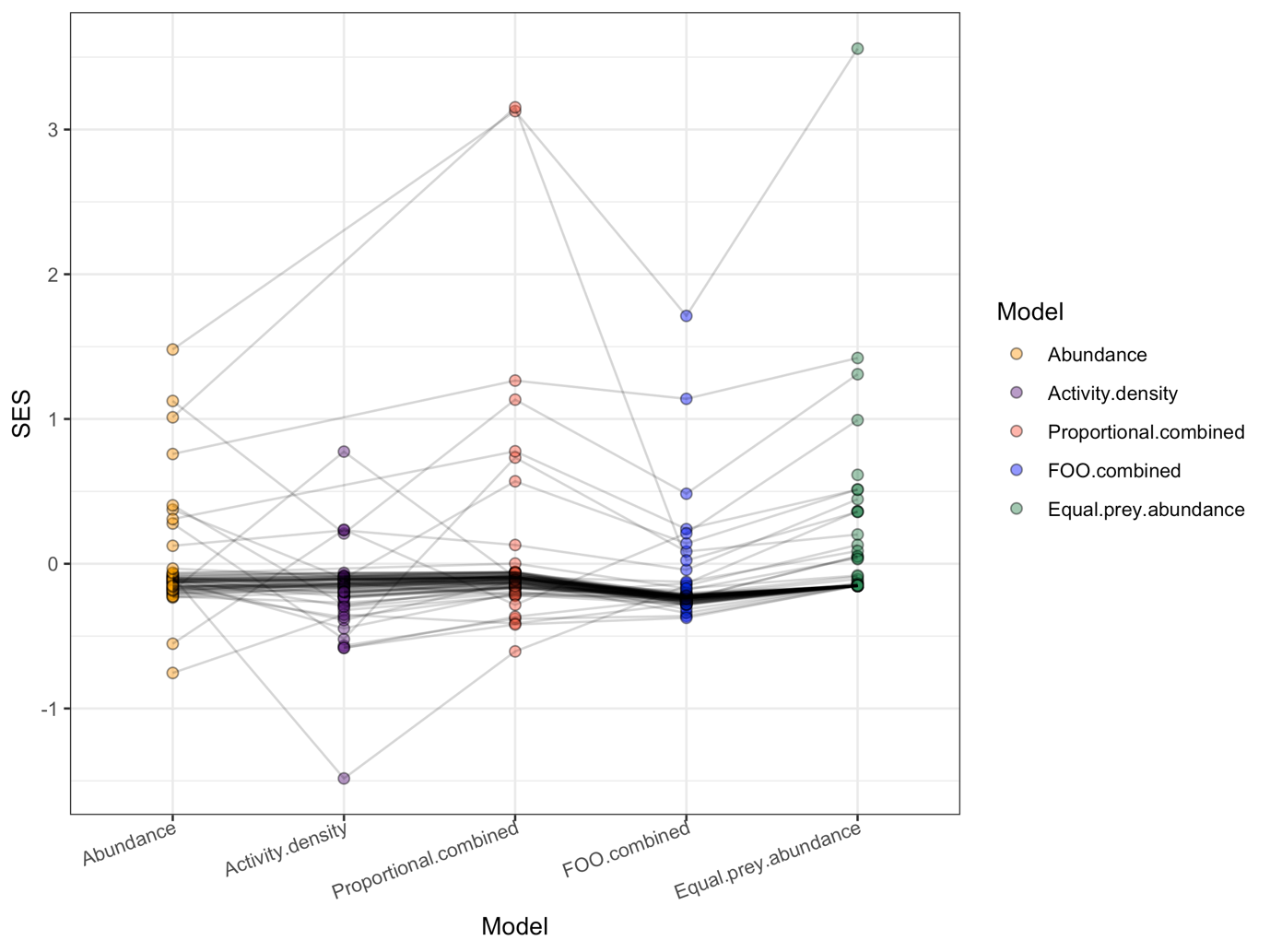

Figure S7: Relative read abundance-based standardised effect sizes (SESs) exported from the comparison of null models against observed interactions. Mean SES values for each prey taxon across the individual consumers were used. Each colour denotes SESs from a different prey choice model. Links are drawn between points representing the same prey taxon. The data very closely resemble those of the presence-absence dietary models, but SESs were generally more positive in the relative read abundance data (Figure S4).

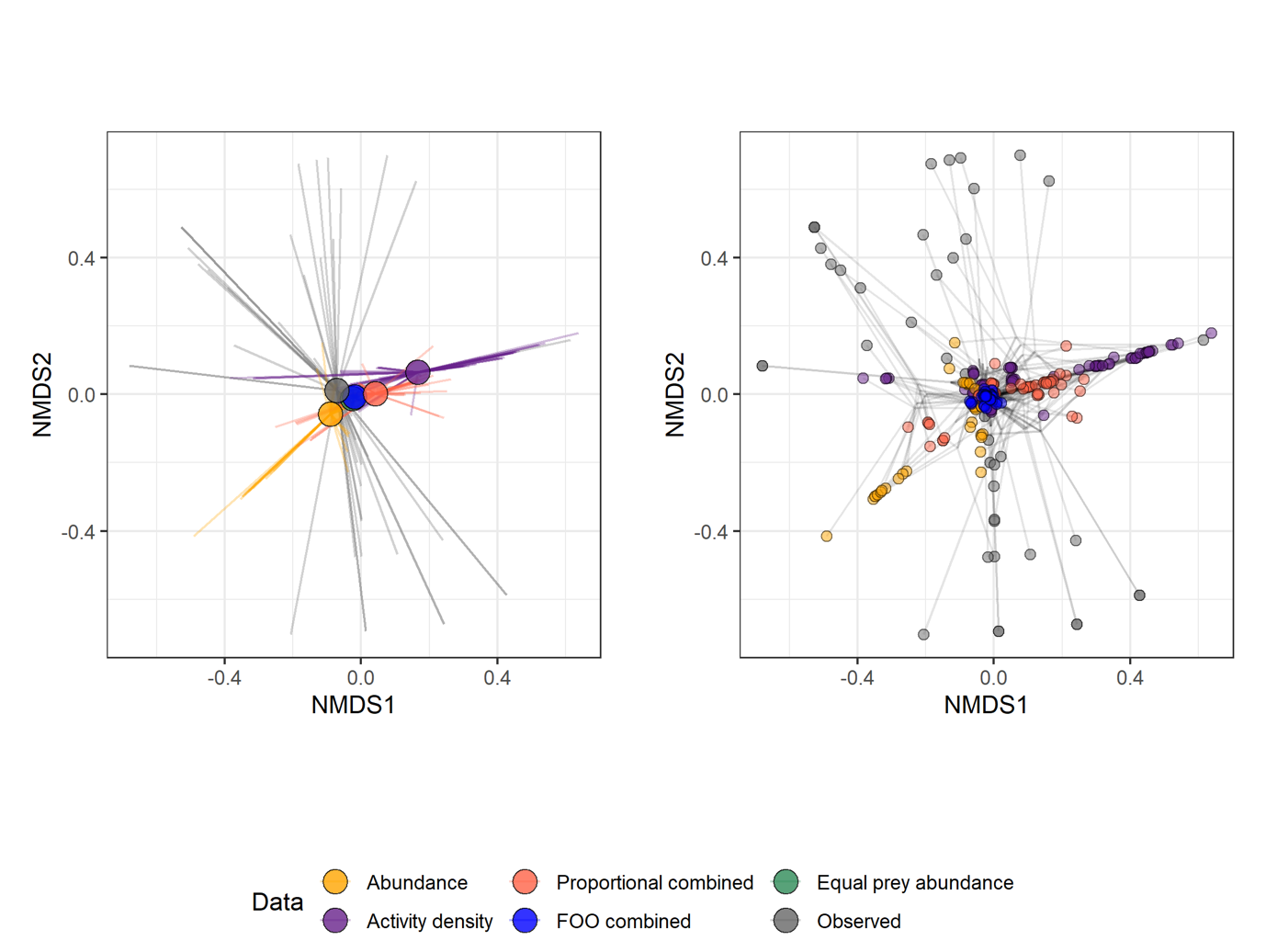

Figure S8: Spider plot derived from non-metric multi-dimensional scaling of observed (relative read abundance-based) and null model-expected spider diets. Colours denote the data source (grey = abundance; other colours are null-model-predicted diets based on the different prey availability datasets). Axes represent a two-dimensional variation in spider diet. In the left plot, the terminal end of each line represents the diet of a spider (whether observed or simulated), joined by the centroids of diets from each data source (larger nodes; mean coordinates in that group), with distance between them indicating their dissimilarity (i.e., proximate points are similar, distant points are dissimilar). In the right plot, each point represents the diet of a spider, with lines joining points representing the same sample and those lines meeting at the mean coordinates for that sample. Mean Euclidean coordinates of the diets, differed in their distance from the mean observed dietary composition based on relative read abundance, with equal prey abundance (Euclidean distance = 0.052), FOO combined (0.055), abundance (0.071), proportional combined (0.115) and activity density (0.244) being progressively further from the observed data, respectively. This relationships between the models and the observed data were similar to those from the presence-absence dietary data, but the equal prey abundance model simulated diets more similar to the observed diets whereas its performance was intermediate from the presence-absence data (Figure 4).

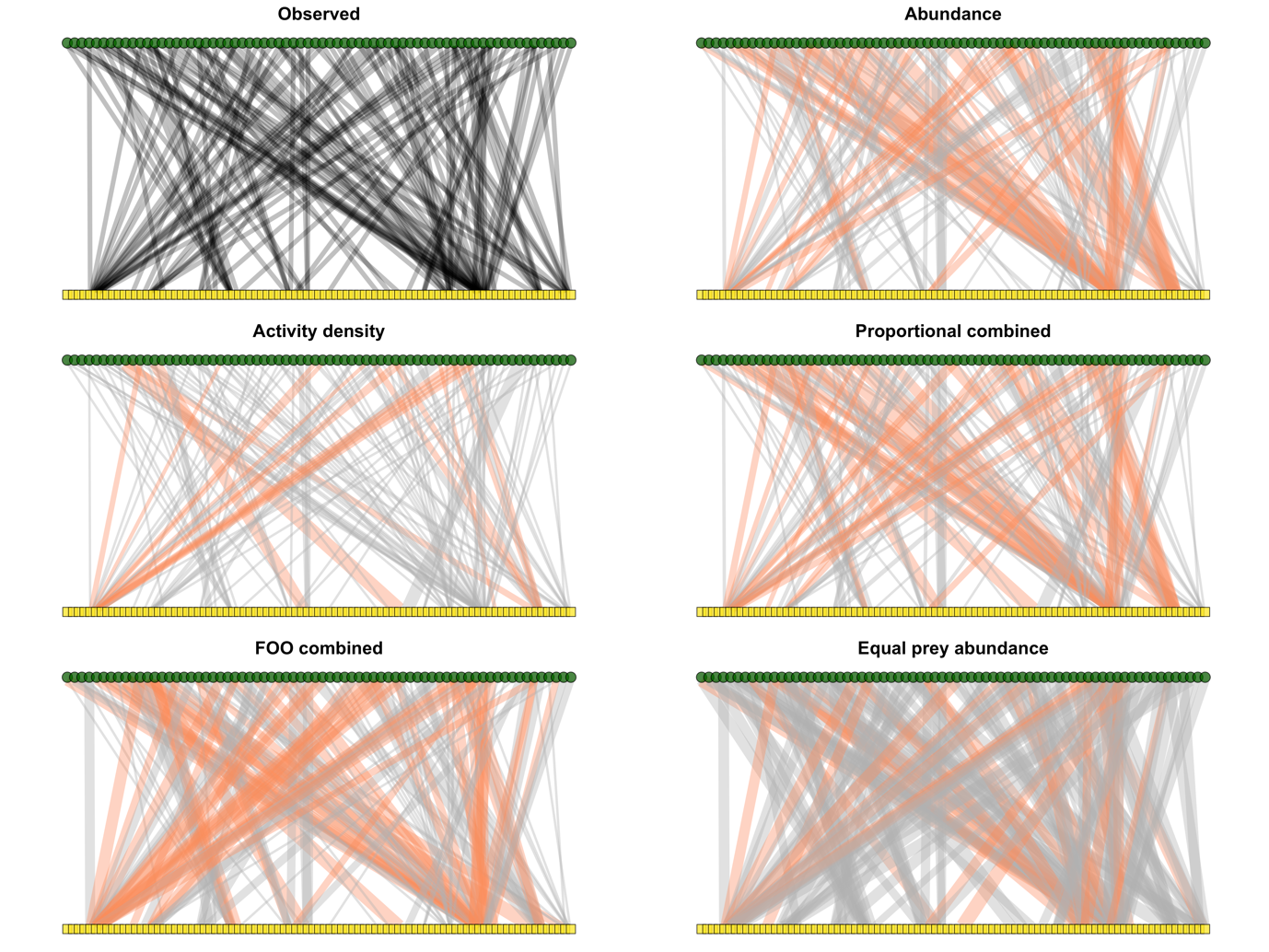

Figure S9: Observed (relative read abundance-based) and null networks produced from the different prey availability datasets. Link weights represent the number of observed interactions for the observed network, and otherwise the frequency of interactions expected based on prey availability according to null models. Red links represent those for which observed interaction frequencies significantly exceeded those expected from null models. No interactions are plotted which were significantly less frequent than expected.

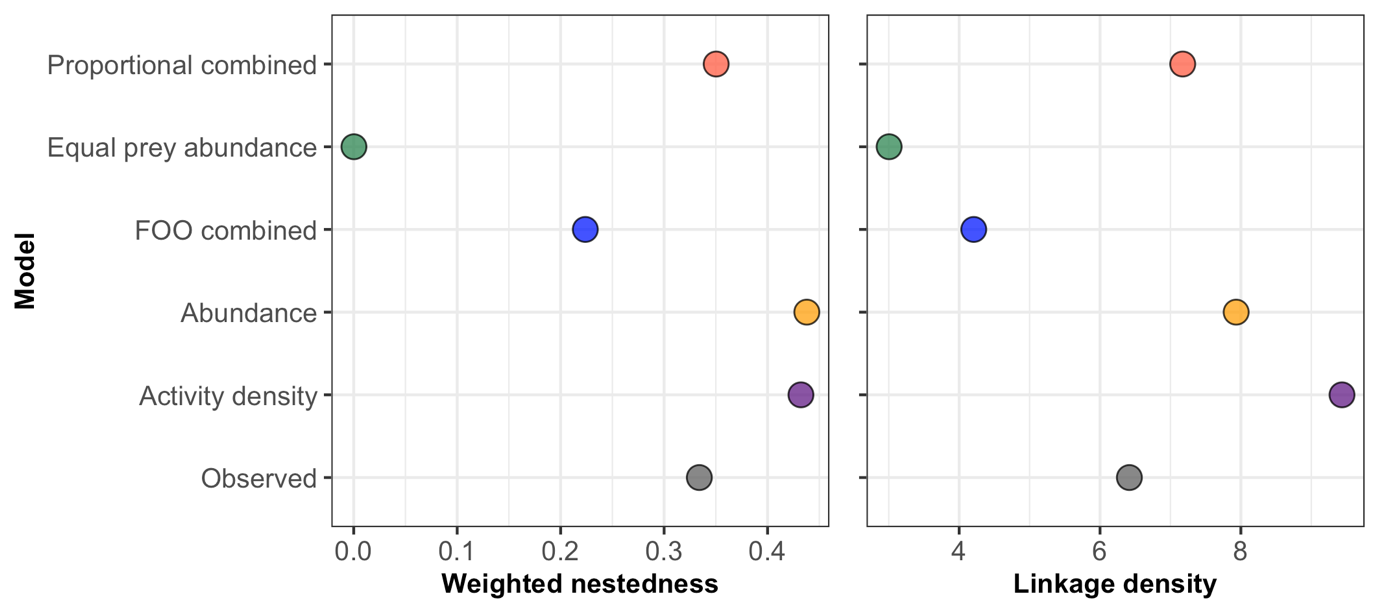

Figure S10: Weighted nestedness and linkage density of the observed (relative read abundance-based) network and the null networks generated from the different prey availability datasets. The observed relative read abundance network has a lower linkage density than the presence-absence network (Figure 6). The null models based on relative read abundance data differ in their relationship to the observed network compared to those of the presence-absence data. The activity density network is marginally more similar to the observed network than the abundance network in terms of nestedness, but the reverse is true of linkage density; both of these relationships are inverted for the presence-absence data.

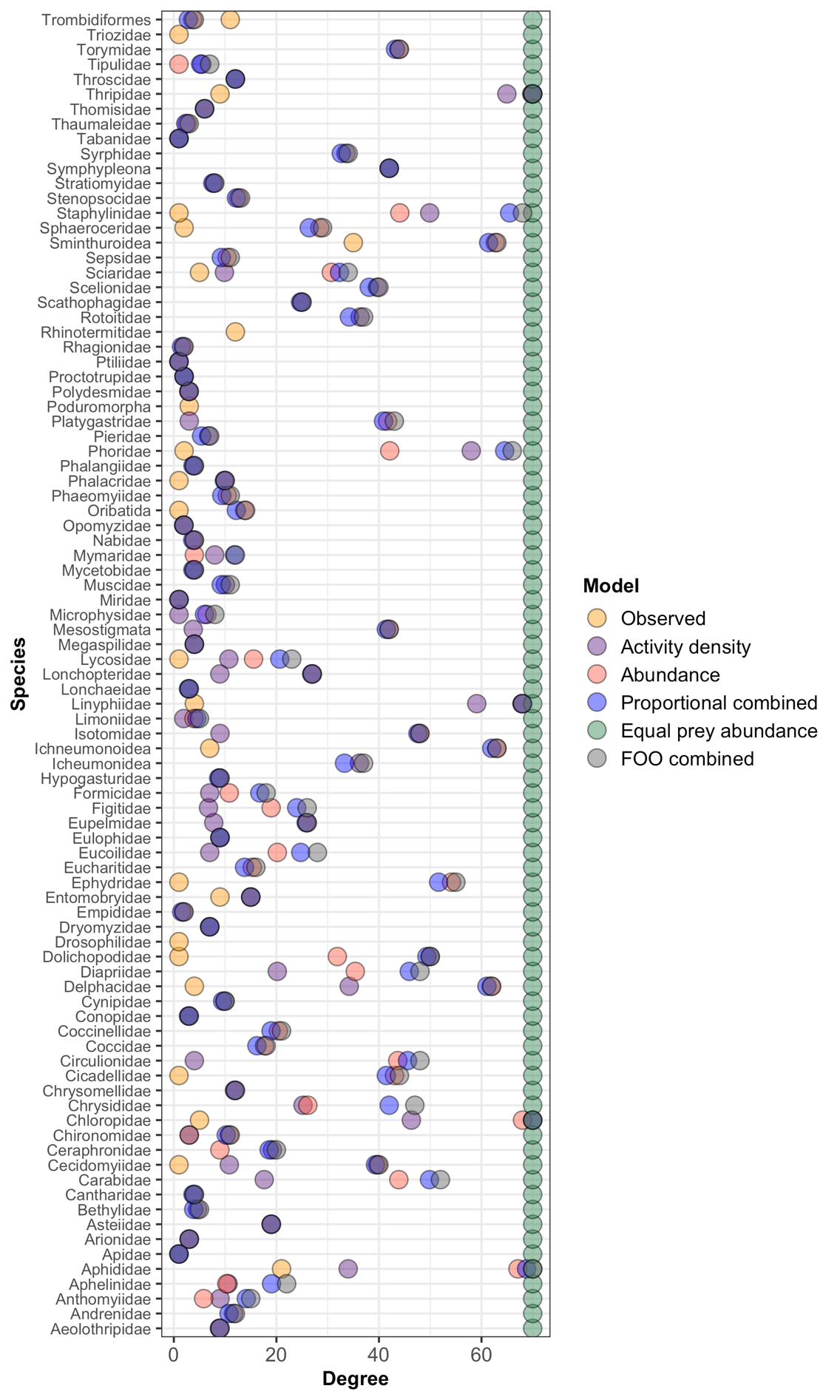

Figure S11: Degree of each node within the observed (relative read abundance-based) and the null networks generated from the different prey availability datasets. The degrees of resource nodes in the relative read abundance-based networks were far more variable and generally much higher, especially compared to the observed network, than those of the presence-absence networks, particularly the degrees of the equal prey abundance null network nodes (Figure S5).

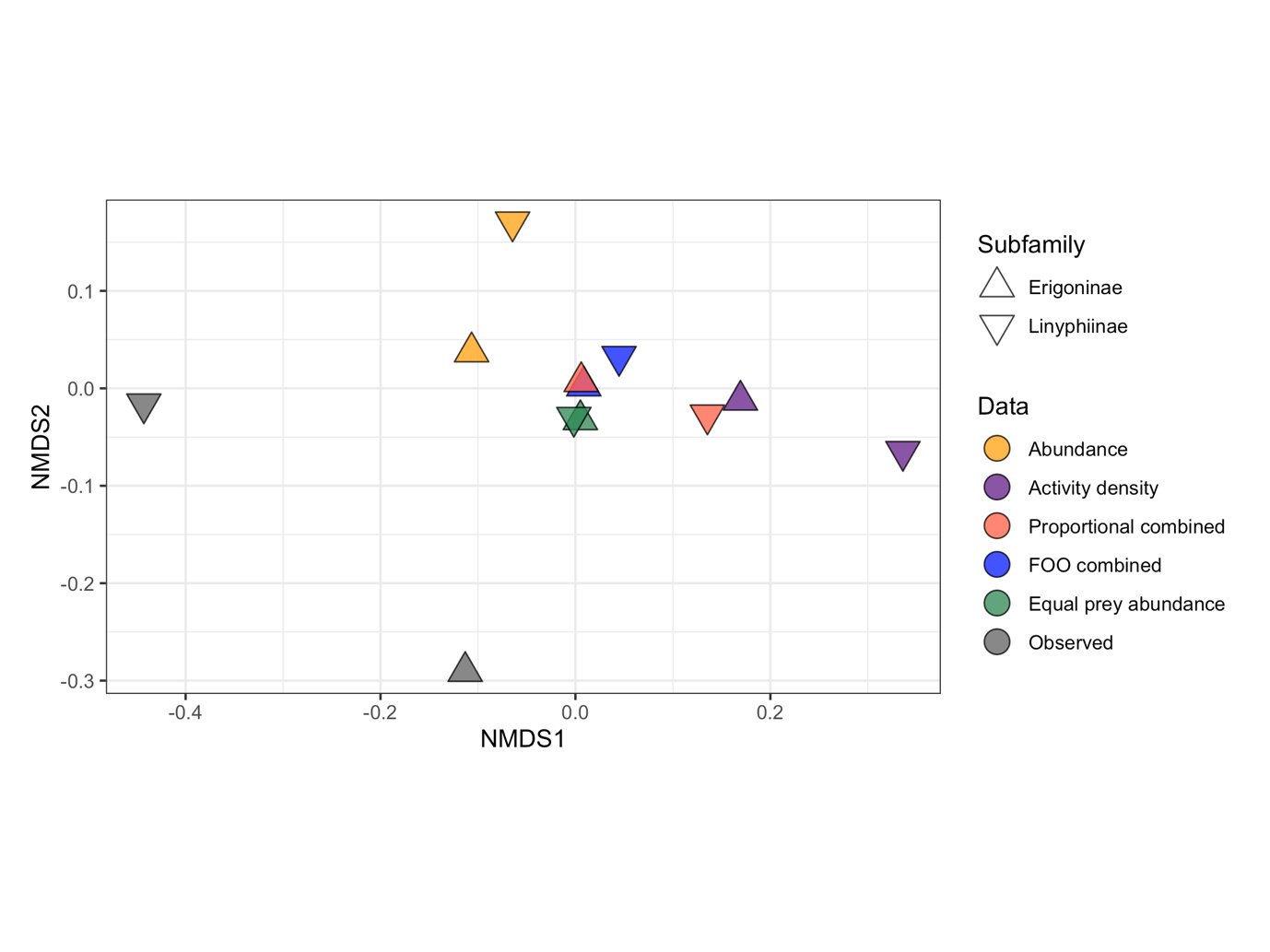

Figure S12: Non-metric multi-dimensional scaling of observed (presence-absence) and null model-expected spider diets for the two subfamilies of spiders. These data were generated in models exactly like those used for Figure 4, but with consumers represented as subfamilies (Linyphiinae and Erigoninae; one Lycosidae individual was removed). Colours denote the data source (grey = abundance; other colours are null-model-predicted diets based on the different prey availability datasets). Axes represent a two-dimensional variation in spider diet. The points are centroids (mean coordinates) of diets from each data source for each subfamily/family with distance between them indicating their dissimilarity (i.e., proximate points are similar, distant points are dissimilar). The diet of both taxa more closely resembled the abundance-based null diets than activity density-based null diets, with activity density null diets being most dissimilar to observed diets for both (Euclidean distance = 0.397 for Erigoninae and 0.780 for Linyphiinae). Erigoninae diets much more closely resembled abundance (Euclidean distance = 0.328) than Linyphiinae (0.422), but this was the most similar null diet for Linyphiinae while Erigoninae diets more closely resembled equal prey abundance (0.284), FOO combined (0.317) and proportional combined (0.320). Replication of each spider group is not, however, even and larger datasets would be required to fully explore this (Linyphiinae = 57, Erigoninae = 13).
